# Early Metabolic Alterations in Cerebrospinal Fluid Fatty Acid Profiles Linked to Cognitive Decline and All-Cause Dementia

**DOI:** 10.1101/2025.08.25.672205

**Authors:** Jacob Apibilla Ayembilla, Yannick N. Wadop, Rebecca Bernal, Biniyam A. Ayele, Murali Sargurupremraj, Xueqiu Jian, Alfred K. Njamnshi, Sudha Seshadri, Alice B. S. Nono Djotsa, Jayandra Jung Himali, Bernard Fongang, Alfred Fonteh, African Initiative on Bioinformatics Online Training in Neurodegenerative Diseases (AI-BOND)

**Author notes:** **Corresponding author** Jacob Apibilla Ayembilla.

## Abstract

**Background:** While All-cause dementia (ACD) may often be characterized by the abnormal deposition of extracellular β-amyloid (Aβ) in the brain cortex and hyperphosphorylated tau (p-tau) as neurofibrillary tangles intracellularly, there is a need to identify early metabolic changes that may accompany these pathological changes.

**Objective:** This study evaluated the predictive value of fatty acids in cerebrospinal fluid (CSF) fractions in differentiating cognitively unimpaired (CU), mild cognitive impairment (MCI), and ACD participants.

**Methods:** CSF fatty acid profiles were analyzed from CU (n=68), MCI (n=38), and ACD (n=37) individuals aged 77.3 ± 7.7 years, sourced from the Huntington Medical Research Institutes (HMRI). Multivariable binary logistic regression identified the most effective CSF fatty acid biomarkers for distinguishing CU, MCI, and ACD groups. The top-performing CSF fatty acid biomarkers were combined with Aβ_42_, tau, and the Aβ_42_/tau ratio to evaluate their collective diagnostic performance. The model was adjusted for covariates, including age, sex, smoking status, hypertension, diabetes, and APOE genotype. Receiver operating characteristic (ROC) curves were generated for the top 10 CSF fatty acids ranked by area under the curve (AUC), sensitivity, and specificity, and their significance was assessed using the DeLong test.

**Results:** The top 10 fatty acids in CSF fractions demonstrated superior discrimination between CU individuals and those with MCI compared to traditional markers such as Aβ_42_, tau, and the Aβ_42_/tau ratio. Furthermore, incorporating a panel of these fatty acid biomarkers alongside Aβ_42_/tau significantly improved diagnostic accuracy. Age, sex, smoking, hypertension, APOE genotype, and diabetes did not significantly influence the model’s performance.

**Conclusion:** This study suggests that changes in fatty acid metabolism occur in early ACD pathology. Thus, strategies that regulate fatty acid metabolism may prevent cognitive decline in an older population.

**Highlights:**

1. This study revealed that CSF fractions’ fatty acids have a better discriminatory power for CU from MCI than CSF Aβ_42_, tau, and Aβ_42_/tau ratio.
2. A panel of CSF fraction fatty acids combined with Aβ_42_/tau remarkably improved its diagnostic performance for differential diagnosis of CU, MCI, and ACD.
3. Clinical evaluation of these fatty acids will strengthen the detection of early cognitive impairment. A prospective large cohort multicenter study of the diagnostic utility of CSF fractions’ fatty acids will provide robust evidence before extensive clinical usage of these fatty acids.

## Introduction

All-cause dementias (ACD) are multifaceted and incurable neurodegenerative conditions, characterized by extensive synapse loss (Chen et al., 2018; Kashyap et al., 2019) cortical atrophy, and neuronal death. These pathological changes lead to progressive cognitive impairment, memory loss, and behavioral disturbances (Castor et al., 2020; Sheng et al., 2012). Central to several ACD pathologies is the abnormal deposition of extracellular β-amyloid (Aβ) plaques in the cerebral cortex and the intracellular aggregation of hyperphosphorylated tau (p-Tau) in neurofibrillary tangles (Davis et al., 2022; Dubois et al., 2018, 2023; Liu et al., 2018), accompanied by functional cholinergic deficits (Richter et al., 2019). Despite advancements in understanding ACD pathogenesis, its precise biochemical underpinnings and the full spectrum of disease-associated biomarkers remain elusive.

Current cerebrospinal fluid (CSF)-based biomarkers, including Aβ_42_, Aβ_40_, total tau (t-tau), phosphorylated tau (p-tau), neurofilament light chain (NfL), glial fibrillary acidic protein (GFAP), and tau phosphorylated at Thr181 (p-tau 181, p-tau217), show promise for early ACD detection (Dubois et al., 2023; Iaccarino et al., 2023; Janelidze et al., 2020). However, the lack of universally accepted diagnostic cutoffs and the influence of comorbidities such as vascular dementia, Lewy body dementia, and traumatic brain injury limit their specificity for AD pathology (Bousiges & Blanc, 2022; Brand et al., 2022; Davidson et al., 2023; Dong et al., 2019; LoBue et al., 2018; Tao et al., 2023). Furthermore, some cognitively unimpaired (CU) individuals exhibit Aβ_42_/tau ratios comparable to those with ACD, reducing the generalizability of this ratio as a differential diagnostic tool (Fonteh et al., 2020; Harrington et al., 2019). These challenges underscore the urgent need for alternative biomarkers that better capture early ACD-specific pathophysiological processes.

Emerging evidence highlights lipid metabolism as a central player in ACD pathology, given its involvement in amyloid precursor protein (APP) processing, inflammation, energy metabolism, membrane remodeling, and blood-brain barrier (BBB) integrity (Cutuli et al., 2022; Di Miceli et al., 2022). Dysregulation of brain lipids, particularly sphingolipids, cholesterol, and glycerophospholipids, exacerbates Aβ aggregation and tau pathology, accelerating ACD progression (Tong et al., 2024). Elevated brain cholesterol levels, for example, promote APP cleavage by β- and γ-secretases, producing Aβ species with varying toxicity levels (Tong et al., 2024).

Despite the recognized importance of lipid dysregulation in ACD, studies investigating CSF lipid alterations as early biomarkers remain sparse. Elevated sphingomyelin levels in prodromal AD (Kosicek et al., 2012), increased phosphocholine, and a reduced lysophosphatidylcholine/phosphatidylcholine (lysoPC/PC) ratio in ACD patients (Walter et al., 2004; Whiley et al., 2014) suggest lipidomic changes may serve as potential biomarkers. Furthermore, fatty acids such as saturated fatty acids (SAFA), monounsaturated fatty acids (MUFA), and polyunsaturated fatty acids (PUFA) are integral to neuronal membranes, yet their metabolic roles in ACD are not fully understood (Fonteh et al., 2014, 2020; Harrington et al., 2009).

The balance between SAFA and UFA impacts APP processing, with high SAFA diets enhancing Aβ deposition and PUFAs, particularly docosahexaenoic acid (DHA, C22:6n-3), showing neuroprotective effects by attenuating glial activation and reducing Aβ burden (Grimm et al., 2011; Oksman et al., 2006). Additionally, DHA and eicosapentaenoic acid (EPA, C20:5n-3) supplementation improve memory, cognitive function, and neuroprotection (Cardoso et al., 2016; Cederholm et al., 2013). However, altered phospholipase A_2_ activity in ACD may lead to phospholipid hydrolysis and PUFA oxidation, generating reactive oxygen species (ROS) and pro-inflammatory mediators, thereby exacerbating ACD pathology. While omega-3 fatty acids are beneficial, omega-6 fatty acids are pro-inflammatory and have been implicated in ACD pathophysiology. Our previous work revealed distinct fatty acid distributions in CSF unesterified fatty acid, supernatant fluid (SF), and brain-derived nanoparticle (NP) fractions. However, their diagnostic utility relative to Aβ and tau biomarkers and the influence of ACD comorbidities remain unexplored. We hypothesize that lipid metabolic changes represent early events in ACD pathogenesis, contributing to aberrant Aβ processing and deposition. Moreover, unesterified fatty acids and fatty acids in the SF and NP fractions may serve as novel biomarkers for distinguishing cognitive unimpaired (CU), mild cognitive impairment (MCI), and ACD.

## Materials and methods

### Data source and participants

This project leverages data collected at the Huntington Medical Research Institutes (HMRI). The study participants were classified as CU, MCI, or ACD based on their medical and neuropsychological diagnoses as described by (Harrington et al., 2013), and they all consented to the study by signing informed consent. They were mainly non-Hispanic whites aged between 70 and 100 years old, comprising 60(41.96%) males and 83(58.04%) females (Table 1). APOE genotyping was performed from peripheral lymphocytes. In this study, we included all study participants with data on demographics, APOE genotype, CSF unesterified, supernatant fluid, and nanoparticles fatty acids, and CSF biomarkers (Aβ_42_, tau, and Aβ_42_/tau ratio) (Harrington et al., 2013). Altogether, 143 subjects were included in this study, made up of 68(47.55%) CU, 38(26.57%) MCI, and 37(25.87%) ACD (Table 1). The ACD was made up of 28 Alzheimer’s disease (AD) and 9 other dementias (OD).

**Table 1:**
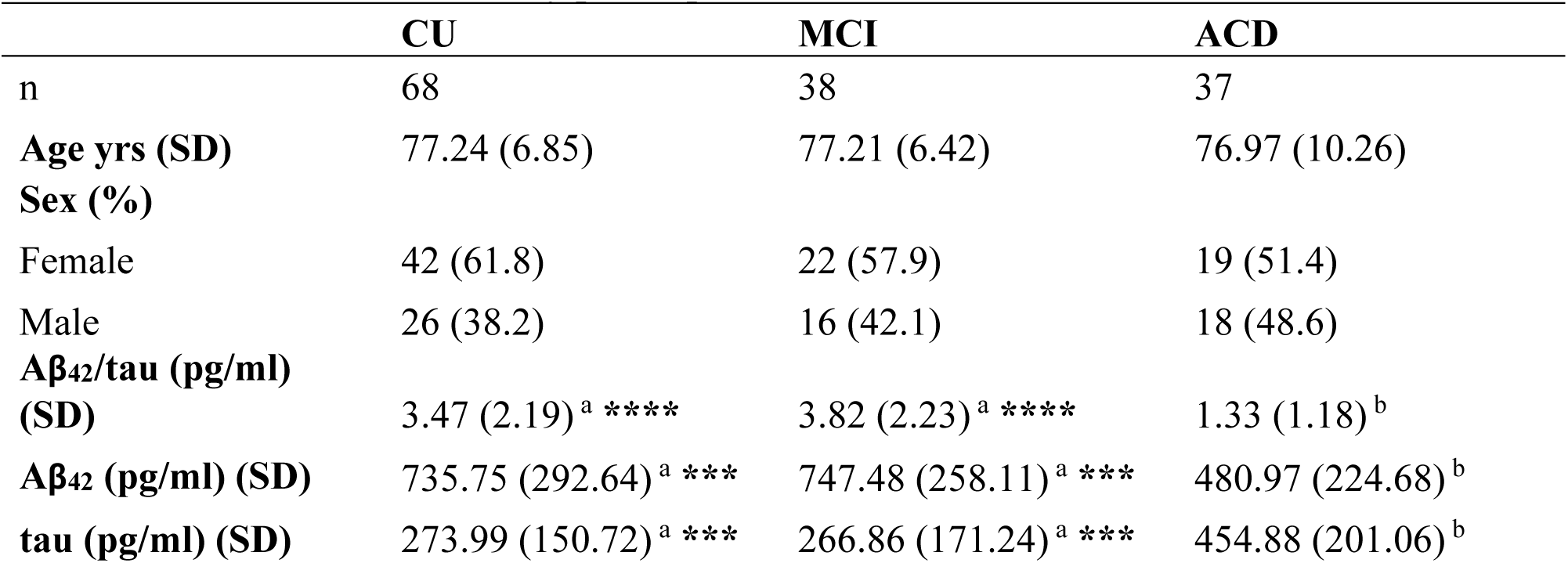
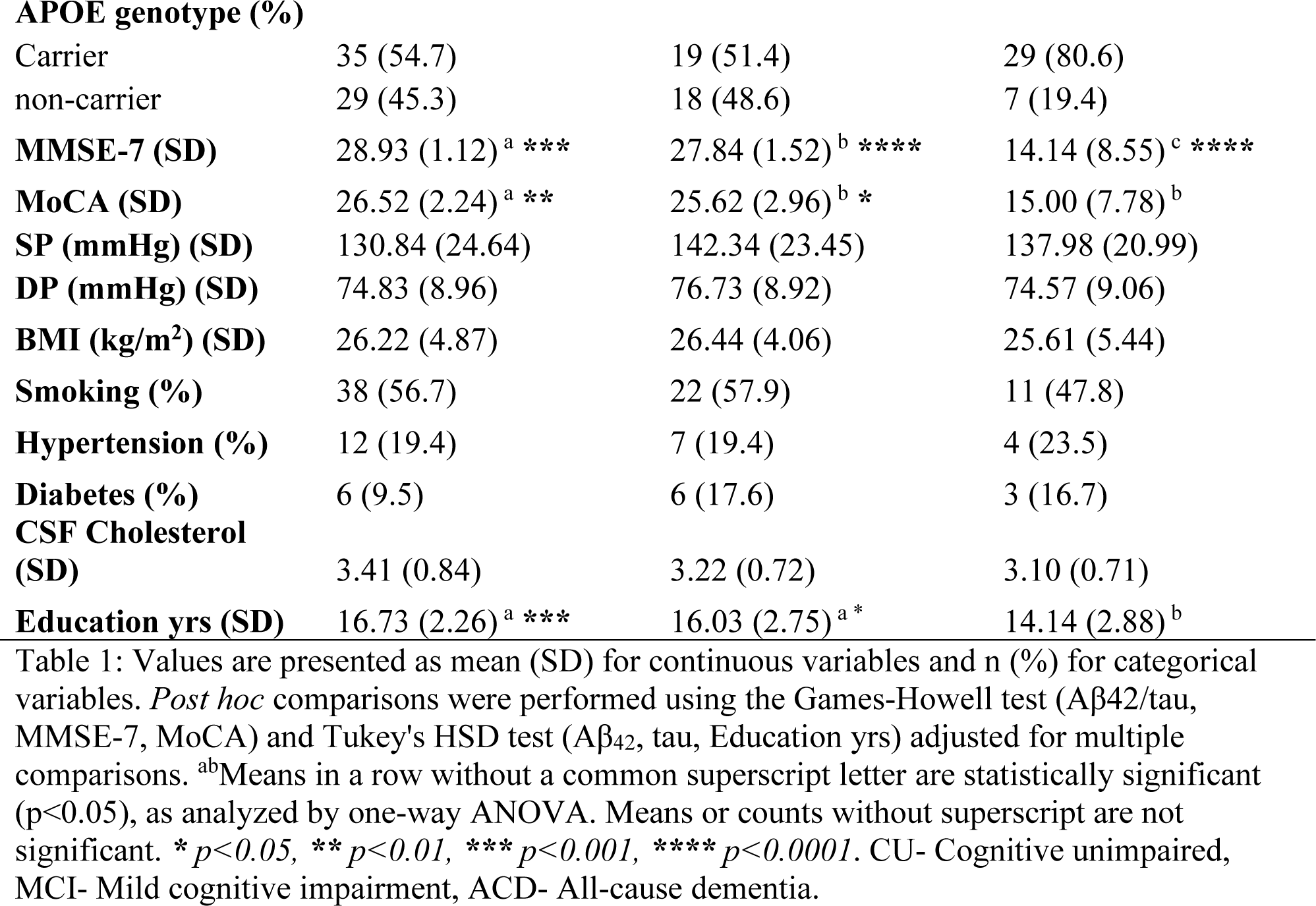
Characteristics of study participants.

### Sample collection and laboratory analysis

The sample collection procedure and laboratory analysis of CSF unesterified, supernatant, and nanoparticles fatty acids and CSF Aβ_42_ and tau have been previously described by (Fonteh et al., 2014). Briefly, after an overnight fast, a lumbar puncture was performed, and approximately 4 mL of CSF sample was collected for analysis. Aliquots of CSF samples were analyzed for Aβ_42_ and tau using a sandwich enzyme-linked immunosorbent assay kit (Innotest b-amyloid (1-42) and Innotest hTAU-Ag, Innogenetics, Gent, Belgium) following the manufacturer’s protocol. CSF aliquots were fractionated by ultracentrifugation, and the supernatant fluid (SF) was harvested, and the pelleted nanoparticles were collected. The fatty acids in each fraction were analyzed with a Gas Chromatography-Mass Spectrometer (GC-MS) (Harrington et al., 2009; Fonteh et al., 2014;) and the GC-MS data obtained were analyzed using Agilent MassHunter Workstation software.

### Statistical analysis

All statistical analyses were performed with R version 4.4.0 (R Core Team, 2021) using the pROC (Turck et al., 2011) and caret package (Kuhn, 2008), and a p-value < 0.05 was considered statistically significant. The continuous variables were reported as mean (SD), and categorical variables were reported as counts (%). Significant differences between the groups were assessed using One-way analysis of variance (ANOVA) or the Kruskal-Wallis test, with a significance threshold of p < 0.05, depending on the data distribution. Post hoc multiple comparisons were tested with Tukey HSD or the Game-Howell test, depending on whether the equal variance assumption was met. Multivariable binary logistic regression analysis with a generalized linear model (glm) was performed to identify the most effective CSF fatty acid biomarkers for distinguishing between CU, MCI, and ACD. The best fitting CSF fatty acids biomarker was integrated with Aβ_42_, tau, and Aβ_42_/tau ratio to assess their impact on the diagnostic performance. Covariates such as age, sex, smoking status, hypertension, diabetes, and APOE genotype were evaluated for their influence on the model. Receiver Operating Characteristics (ROC) curves were generated for the top 10 CSF fractions’ fatty acid biomarkers based on the area under the curve (AUC), sensitivity, and specificity. The DeLong test was performed to identify the significant biomarkers or panels.

## Results

### Characteristics of the study participants

### Diagnostic Performance of CSF Unesterified Fatty Acids in Differentiating CU, MCI, and ACD

The diagnostic performance of the biomarkers studied was evaluated using multivariable logistic regression and ROC analyses (**Table S1**; **Fig. 1**).

**Fig. 1.**
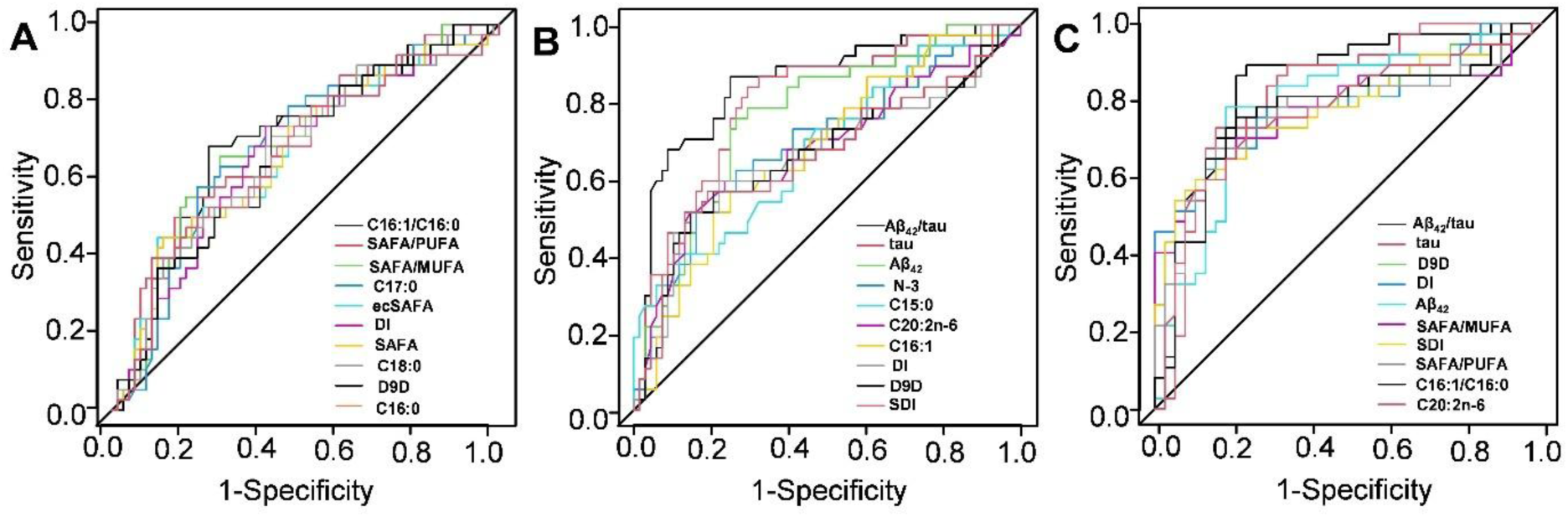
ROC curve analysis of the best performing CSF unesterified fatty acids in discriminating (A) CU from MCI, (B) CU from ACD, and (C) MCI from ACD. Where DI- Desaturates index, ecSAFA- even chain saturated fatty acids, D9D- Delta-9 desaturase, SDI- Saturation Desaturates Index.

### Discrimination Between CU and MCI

CSF unesterified fatty acids outperformed canonical biomarkers (Aβ_42_, tau, and Aβ_42_/tau) in differentiating CU individuals from those with MCI. The unesterified fatty acids demonstrated AUC values ranging from 0.66 to 0.70, sensitivity between 0.39 and 0.76, and specificity between 0.57 and 0.88. In comparison, Aβ_42_, tau, and Aβ_42_/tau recorded AUCs of 0.53, 0.54, and 0.55, respectively, with sensitivity ranging from 0.76 to 0.82 and specificity ranging from 0.32 to 0.81 (Table S1, Fig. 1A).

The ratio of unesterified palmitoleic acid to palmitic acid (C16:1/C6:0) exhibited the highest diagnostic accuracy, with an AUC of 0.70, sensitivity of 0.68, and specificity of 0.75. Other notable ratios included unesterified saturated to unesterified polyunsaturated fatty acids (SAFA/PUFA), unesterified saturated to monounsaturated fatty acids (SAFA/MUFA), and unesterified unsaturated to saturated fatty acids (DI), which achieved AUCs of 0.69 with comparable sensitivity (0.55–0.74) and specificity (0.60–0.81). The lowest performer among the top 10 biomarkers was palmitic acid (unesterified C16:0), with an AUC of 0.66, sensitivity of 0.39, and specificity of 0.90 (Table S1).

Statistical analysis revealed significant differences between Aβ_42_ and the top-performing fatty acids (e.g., unesterified C16:1/C16:0, unesterified SAFA/PUFA), as well as tau, at a p-value threshold of <0.05 (Supplementary Table S2).

### Discrimination Between MCI and ACD

In distinguishing MCI from ACD, the canonical biomarkers (Aβ_42_, tau, and Aβ_42_/tau) demonstrated superior performance with AUCs of 0.79, 0.82, and 0.85, respectively. CSF unesterified fatty acids exhibited slightly lower diagnostic performance, with AUCs ranging from 0.77 to 0.79, sensitivity between 0.65 and 0.78, and specificity between 0.71 and 0.87 (Table S1, Fig. 1B).

Among the fatty acids, total desaturase 9 activity (Free D9D) and the ratio of unesterified unsaturated to saturated fatty acids (DI) both achieved an AUC of 0.79, matching Aβ_42_. Lower-performing biomarkers included unesterified palmitoleic acid/palmitic acid ratio (C16:1/C16:0), total saturated to polyunsaturated fatty acids ratio (SAFA/PUFA), and unesterified eicosadienoic acid (C20:2n-6), with AUCs of 0.77 (Table S1). There were no statistically significant differences between the AUCs of Aβ_42_, tau, and Aβ_42_/tau and the top-performing fatty acids (Supplementary Table S2).

### Discrimination Between CU and ACD

The canonical biomarkers again showed the highest diagnostic accuracy, with AUCs of 0.76, 0.80, and 0.85 for Aβ_42_, tau, and Aβ_42_/tau, respectively. Sensitivity ranged from 0.76 to 0.87, while specificity ranged from 0.71 to 0.75 (Table S1; Fig. 1)

Among CSF unesterified fatty acids, omega-3 fatty acids (ω-3) achieved the highest AUC (0.69), followed by pentadecanoic acid (C15:0) with an AUC of 0.68. The unesterified stearic acid/oleic acid ratio (C18:1/C18:0) (SDI) recorded the lowest AUC (0.66) (Table S2; Fig. 1C). Significant differences were observed between the AUCs of Aβ_42_/tau and the top fatty acids, but no differences were found between Aβ_42_, tau, and the fatty acids (Supplementary Table S1).

### Combined Biomarker Performance

To assess the additive value of CSF unesterified fatty acids, they were combined with Aβ_42_/tau. This integration improved the diagnostic performance across all group comparisons:

- **CU vs. MCI**: Aβ_42_/tau alone had an AUC of 0.55, which increased to 0.70–0.71 with fatty acids. The unesterified SAFA/PUFA ratio provided the greatest improvement (Table S3).
- **MCI vs. ACD**: The AUC of Aβ_42_/tau increased from 0.85 to 0.86–0.93 with the addition of fatty acids, with unesterified eicosadienoic acid (C20:2(n-6)) showing the best enhancement (Table S3).
- **CU vs. ACD**: The AUC of Aβ_42_/tau increased from 0.85 to 0.85–0.87. Free omega-3 fatty acids (N-3) and pentadecanoic acid (C15:0) were the most effective enhancers (Table S3).

Analysis of covariates showed no significant impact on the combined models (Table 2).

**Table 2:** Effect of covariates on the best CSF unesterified fatty acids model for discriminating CU, MCI, and AD.

| Model | n | AUC (95% CI) | Sensitivity | Specificity |
| --- | --- | --- | --- | --- |
| **CU versus MCI** |  |  |  |  |
| $A\beta_{42}/\tau + C17:0$ | 68 vs 38 | 0.74(0.63-0.84) | 0.85 | 0.59 |
| $A\beta_{42}/\tau + C17:0 + \text{Age}$ | 68 vs 38 | 0.74(0.64-0.85) | 0.71 | 0.75 |
| $A\beta_{42}/\tau + C17:0 + \text{Sex}$ | 68 vs 38 | 0.76(0.65-0.87) | 0.74 | 0.82 |
| $A\beta_{42}/\tau + C17:0 + \text{Diabetes}$ | 68 vs 38 | 0.75(0.65-0.86) | 0.71 | 0.75 |
| $A\beta_{42}/\tau + C17:0 + \text{Hypertension}$ | 68 vs 38 | 0.74(0.63-0.84) | 0.85 | 0.59 |
| Aβ42/tau + C17:0 + APOE | 68 vs 38 | 0.74(0.63-0.84) | 0.85 | 0.59 |
| Aβ42/tau + C17:0 + Smoking | 68 vs 38 | 0.74(0.63-0.84) | 0.85 | 0.59 |
| **MCI versus ACD** |  |  |  |  |
| Aβ42/tau + C20:2n-6 | 38 vs 37 | 0.84(0.71-0.97) | 0.89 | 0.76 |
| Aβ42/tau + C20:2n-6 + Age | 38 vs 37 | 0.85(0.72-0.98) | 0.89 | 0.76 |
| Aβ42/tau + C20:2n-6 + Sex | 38 vs 37 | 0.85(0.72-0.97) | 0.89 | 0.76 |
| Aβ42/tau + C20:2n-6 + Diabetes | 38 vs 37 | 0.84(0.72-0.97) | 0.89 | 0.76 |
| Aβ42/tau + C20:2n-6 + Hypertension | 38 vs 37 | 0.84(0.71-0.97) | 0.89 | 0.76 |
| Aβ42/tau + C20:2n-6 + APOE | 38 vs 37 | 0.84(0.71-0.97) | 0.89 | 0.76 |
| Aβ42/tau + C20:2n-6 + Smoking | 38 vs 37 | 0.84(0.71-0.97) | 0.89 | 0.76 |
| **CU versus ACD** |  |  |  |  |
| Aβ42/tau + C16:1 | 68 vs 37 | 0.86(0.73-0.98) | 0.78 | 0.89 |
| Aβ42/tau + C16:1 + Age | 68 vs 37 | 0.87(0.74-0.99) | 0.72 | 0.98 |
| Aβ42/tau + C16:1 + Sex | 68 vs 37 | 0.86(0.73-0.99) | 0.78 | 0.89 |
| Aβ42/tau + C16:1 + Diabetes | 68 vs 37 | 0.84(0.72-0.97) | 0.78 | 0.84 |
| Aβ42/tau + C16:1 + Hypertension | 68 vs 37 | 0.86(0.73-0.98) | 0.78 | 0.89 |
| Aβ42/tau + C16:1 + APOE | 68 vs 37 | 0.86(0.73-0.98) | 0.78 | 0.89 |
| Aβ42/tau + C16:1 + Smoking | 68 vs 37 | 0.86(0.73-0.98) | 0.78 | 0.89 |

### Diagnostic Performance of CSF Supernatant Fluid Fatty Acids in Differentiating CU, MCI, and ACD

The diagnostic utility of CSF supernatant fluid fatty acids for distinguishing CU, MCI, and ACD was assessed using multivariable binary logistic regression and ROC curve analysis (Table S4, Fig. 2).

**Fig. 2.**
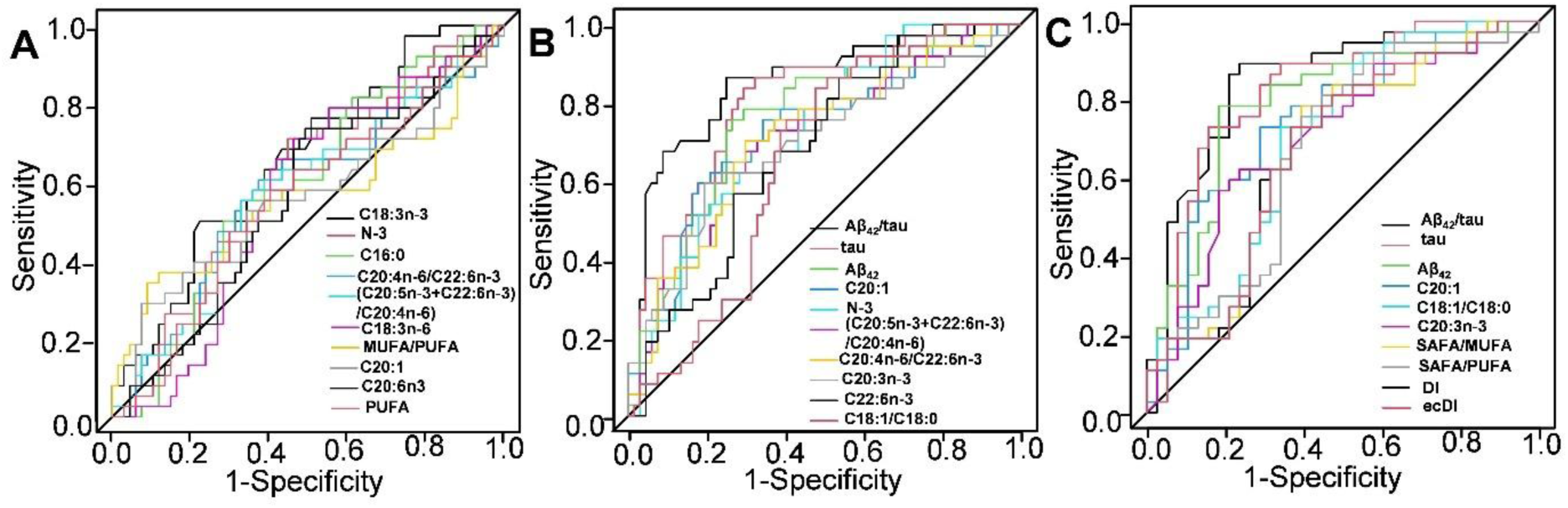
ROC curve analysis of the best performing CSF supernatant fluid fatty acids and CSF biomarkers in discriminating (A) CU vs. MCI, (B) CU vs. ACD, and (C) MCI vs. ACD. Where DI- Desaturates index, SDI- Saturation Desaturates Index, ecDI- even chain Desaturates Index.

### CU vs. MCI

The top 10 supernatant fluid fatty acids emerged as superior biomarkers compared to Aβ, tau, and the Aβ/tau ratio, with AUC values ranging from 0.56 to 0.66. Sensitivity and specificity for these fatty acids varied from 0.29–0.74 and 0.51–0.93, respectively, outperforming Aβ (AUC = 0.53, sensitivity = 0.82, specificity = 0.32), tau (AUC = 0.54, sensitivity = 0.34, specificity = 0.81), and Aβ/tau (AUC = 0.55, sensitivity = 0.76, specificity = 0.44) (Table S4). Among these biomarkers, alpha-linolenic acid (ALA, C18:3(n-3)) demonstrated the highest diagnostic accuracy (AUC = 0.66, sensitivity = 0.50, specificity = 0.78), while total polyunsaturated fatty acids (PUFAs) had the lowest AUC (0.56). A statistically significant difference was observed between the AUC of Aβ_42_ and ALA (Supplementary Table S5). The ROC curves for CU vs. MCI are presented in Fig. 2A.

### MCI vs. ACD

Supernatant fluid Aβ, tau, and the Aβ/tau ratio showed strong classification performance, with AUC values of 0.79, 0.82, and 0.84, respectively. The top-performing fatty acids exhibited AUC values between 0.67 and 0.76, with sensitivity ranging from 0.62–0.81 and specificity from 0.63–0.76 (Table S4). Eicosenoic acid (C20:1) had the highest AUC (0.76), sensitivity (0.73), and specificity (0.71), while the total even chain unsaturated fatty acids/total even chain saturated fatty acids ratio (ecDI) had the lowest AUC (0.67). Aβ/tau demonstrated significantly better performance than several fatty acid biomarkers, including SAFA/MUFA, SAFA/PUFA, and ecDI (p < 0.05, Supplementary Table S5). No significant differences were found between Aβ_42_, tau, and the top-performing fatty acids. Fig. 2B shows the ROC curves for MCI vs. ACD.

### CU vs. ACD

Aβ_42_, tau, and the Aβ/tau ratio were the strongest classifiers, with AUC values of 0.76, 0.80, and 0.86, sensitivities of 0.76, 0.84, and 0.87, and specificities of 0.71, 0.74, and 0.75, respectively. The top fatty acids yielded AUC values of 0.66–0.73, sensitivities of 0.59–0.89, and specificities of 0.46–0.82 (Table S4). Eicosenoic acid (C20:1) recorded the highest AUC (0.73), sensitivity (0.59), and specificity (0.82), followed closely by total N-3 fatty acids (AUC = 0.72, sensitivity = 0.89, specificity = 0.45). The stearic acid/oleic acid ratio (C18:1/C18:0) (SDI) had the lowest AUC (0.66), sensitivity (0.84), and specificity (0.52). Significant differences were noted between tau and C18:1/C18:0, Aβ_42_/tau, and fatty acids such as C22:6n-3, C20:3(n-3), and (C20:5n-3+C22:6n-3)/C20:4n-6 (p < 0.05, Supplementary Table S5). Fig. 2C illustrates the ROC curves for CU vs. ACD.

### Enhancement of Aβ/tau Performance by Fatty Acids

Combining supernatant fluid fatty acids with Aβ/tau improved diagnostic performance across all comparisons. For CU vs. MCI, the AUC of Aβ/tau increased from 0.55 to a range of 0.60–0.67, with ALA (C18:3n-3) providing the greatest improvement (AUC = 0.67). For MCI vs. ACD, the addition of C20:3n-3) and C20:1 to Aβ/tau enhanced the AUC from 0.85 to 0.94, while C18:1/C18:0 (SDI) had the least impact (AUC = 0.90). Similarly, for CU vs. ACD, the AUC of Aβ/tau ratio improved from 0.85 to 0.88–0.92 with the addition of C20:1 and N-3 (Table S6). The addition of covariates had no statistically significant effect on the best model (Table 3).

**Table 3:** Effect of covariates on the best model supernatant fluid fatty acids for this discriminating CU, MCI, and ACD.

| Variable | n | AUC (95% CI) | Sensitivity | Specificity |
| --- | --- | --- | --- | --- |
| **CU vs. MCI** |  |  |  |  |
| $A\beta_{42}$/tau + C18:3n-3 | 68 vs 38 | 0.70(0.59-0.81) | 0.62 | 0.71 |
| $A\beta_{42}$/tau + C18:3n-3 + Age | 68 vs 38 | 0.70(0.59-0.81) | 0.62 | 0.7 |
| $A\beta_{42}$/tau + C18:3n-3 + Sex | 68 vs 38 | 0.70(0.60-0.81) | 0.68 | 0.68 |
| $A\beta_{42}$/tau + C18:3n-3 + Smoking | 68 vs 38 | 0.70(0.59-0.81) | 0.88 | 0.44 |
| $A\beta_{42}$/tau + C18:3n-3 + Hypertension. | 68 vs 38 | 0.70(0.59-0.80) | 0.65 | 0.7 |
| $A\beta_{42}$/tau + C18:3n-3 + Diabetes | 68 vs 38 | 0.72(0.62-0.82) | 0.71 | 0.64 |
| $A\beta_{42}$/tau + C18:3n-3 + APOE genotype. | 68 vs 38 | 0.70(0.59-0.81) | 0.62 | 0.7 |
| **MCI vs. ACD** |  |  |  |  |
| $A\beta_{42}$/tau + C20:3n-3 + C20:1 | 38 vs 37 | 0.94(0.89-1.00) | 0.97 | 0.82 |
| $A\beta_{42}$/tau + C20:3n-3 + C20:1 + Age | 38 vs 37 | 0.94(0.88-1.00) | 0.97 | 0.82 |
| $A\beta_{42}$/tau + C20:3n-3 + C20:1 + Sex | 38 vs 37 | 0.94(0.89-1.00) | 0.97 | 0.82 |
| $A\beta_{42}$/tau + C20:3n-3 + C20:1 + Smoking | 38 vs 37 | 0.93(0.86-1.00) | 1 | 0.76 |
| $A\beta_{42}$/tau + C20:3n-3 + C20:1 + Hypertension | 38 vs 37 | 0.92(0.84-1.00) | 0.79 | 0.91 |
| Aβ<sub>42</sub>/tau + C20:3n-3 + C20:1 + Diabetes | 38 vs 37 | 0.91(0.83-1.00) | 0.93 | 0.89 |
| Aβ<sub>42</sub>/tau + C20:3n-3 + C20:1 + APOE genotype | 38 vs 37 | 0.92(0.84-1.00) | 0.79 | 0.94 |
| **CU vs. ACD** |  |  |  |  |
| Aβ<sub>42</sub>/tau + C20:1 + ω-3 | 68 vs 37 | 0.92(0.86-0.98) | 0.81 | 0.91 |
| Aβ<sub>42</sub>/tau + C20:1 + ω-3 + Age | 68 vs 37 | 0.92(0.86-0.98) | 0.87 | 0.87 |
| Aβ<sub>42</sub>/tau + C20:1 + ω-3 + Sex | 68 vs 37 | 0.92(0.86-0.98) | 0.84 | 0.88 |
| Aβ<sub>42</sub>/tau + C20:1 + ω-3 + Smoking | 68 vs 37 | 0.89(0.78-1.00) | 0.82 | 0.92 |
| Aβ<sub>42</sub>/tau + C20:1 + ω-3 + Hypertension | 68 vs 37 | 0.90(0.79-1.00) | 0.88 | 0.92 |
| Aβ<sub>42</sub>/tau + C20:1 + ω-3 + Diabetes | 68 vs 37 | 0.89(0.79-1.00) | 0.82 | 0.9 |
| Aβ<sub>42</sub>/tau + C20:1 + ω-3 + APOE genotype | 68 vs 37 | 0.91(0.81-1.00) | 0.82 | 0.95 |

### Diagnostic Performance of CSF Nanoparticle Fatty Acids in Differentiating CU, MCI, and ACD

To assess their diagnostic utility, the top 10 CSF nanoparticle fatty acid biomarkers were evaluated for their ability to distinguish CU individuals, MCI patients, and those with ACD (Table S7, Fig.3).

**Fig. 3.**
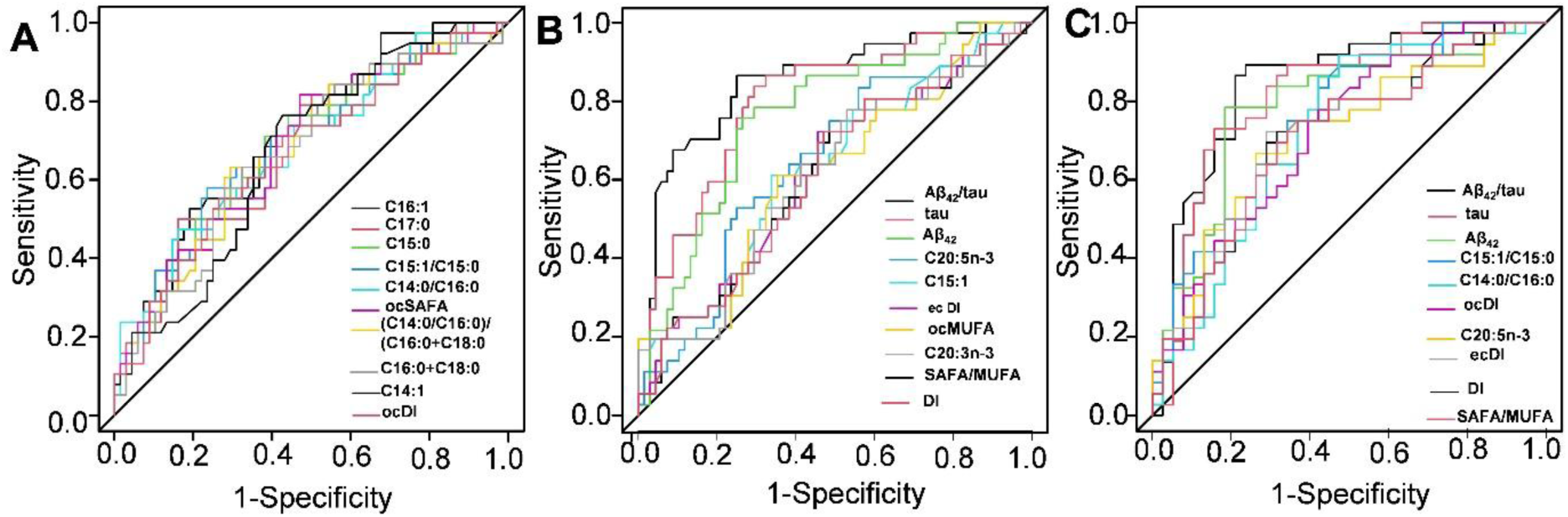
ROC curve analysis of the best performing CSF nanoparticles fatty acids in discriminating (A) CU vs. MCI, (B) CU vs. ACD, and (C) MCI vs. ACD. Where DI- Desaturates index, ecDI- even chain Desaturates Index, ocDI- odd chain Desaturates Index, ocSAFA- odd chain saturated fatty acids, ocMUFA- odd chain Monounsaturated fatty acids.

### CU vs. MCI

The fatty acid biomarkers demonstrated superior diagnostic performance compared to traditional biomarkers (Aβ, tau, and Aβ/tau ratio). The nanoparticle fatty acids achieved AUCs ranging from 0.67 to 0.71, sensitivities of 0.50–0.82, and specificities of 0.57–0.84, outperforming Aβ, tau, and Aβ/tau ratio (AUCs: 0.53–0.55; sensitivities: 0.34–0.82; specificities: 0.32–0.81) (Table S7). Among the fatty acids, palmitoleic acid (C16:1) exhibited the highest performance with an AUC of 0.71, sensitivity of 0.53, and specificity of 0.81. Other high-performing fatty acids included heptadecanoic acid (C17:0), pentadecanoic acid (C15:0), and the pentadecenoic acid/pentadecanoic acid ratio (C15:1/C15:0), all with AUCs of 0.69. ROC curves for these biomarkers are shown in Fig. 3A. Statistically significant differences were observed between Aβ_42_ and several fatty acid biomarkers, including palmitoleic acid, pentadecanoic acid, and heptadecanoic acid (Supplementary Table S8).

### MCI vs. ACD

Traditional biomarkers (Aβ, tau, and Aβ/tau ratio) were the most effective for differentiating MCI from ACD, with AUCs of 0.79, 0.82, and 0.85, respectively (Table S7). The top-performing nanoparticle fatty acids had AUCs ranging from 0.71 to 0.75, sensitivities of 0.67–0.92, and specificities of 0.53–0.74. Pentadecenoic acid/pentadecanoic acid ratio (C15:1/C15:0) recorded the highest AUC of 0.75, while total unsaturated fatty acids/total saturated fatty acids ratio (DI) and total odd unsaturated fatty acids/total odd saturated fatty acids ratio (ocDI) had the lowest AUCs (0.71) (Table S7). Statistically significant differences were observed between Aβ/tau and several fatty acid biomarkers (Supplementary Table S8). ROC curves for MCI vs. ACD are shown in Fig. 3B.

### CU vs. ACD

Aβ, tau, and Aβ/tau ratio were the best-performing biomarkers for distinguishing CU from ACD, with AUCs of 0.76, 0.80, and 0.85, respectively (Table S7). Among fatty acids, eicosapentaenoic acid (EPA, C20:5(n-3) showed the highest diagnostic performance with an AUC of 0.65, sensitivity of 0.53, and specificity of 0.75. Other fatty acids, including total odd monounsaturated fatty acids (ocMUFA) and **e**icosatrienoic acid (C20:3(n-3), showed lower AUCs (0.61). Except for EPA, the differences between Aβ_42_ and the fatty acid biomarkers were statistically significant (Supplementary Table S8). ROC curves for CU vs. ACD are presented in Fig. 3C.

### Enhancing Diagnostic Accuracy

Combining fatty acids with traditional biomarkers improved diagnostic performance. For CU vs. MCI, adding palmitoleic acid improved the AUC of the Aβ_42_/tau ratio from 0.55 to 0.71. For MCI vs. ACD, eicosapentaenoic acid increased the Aβ_42_/tau ratio AUC from 0.85 to 0.89. Similarly, for CU vs. ACD, eicosapentaenoic acid and pentadecanoic acid enhanced the Aβ_42_ AUC from 0.85 to 0.86 (Table S9). The inclusion of covariates in the diagnostic models did not yield statistically significant differences (Table 4).

**Table 4:**
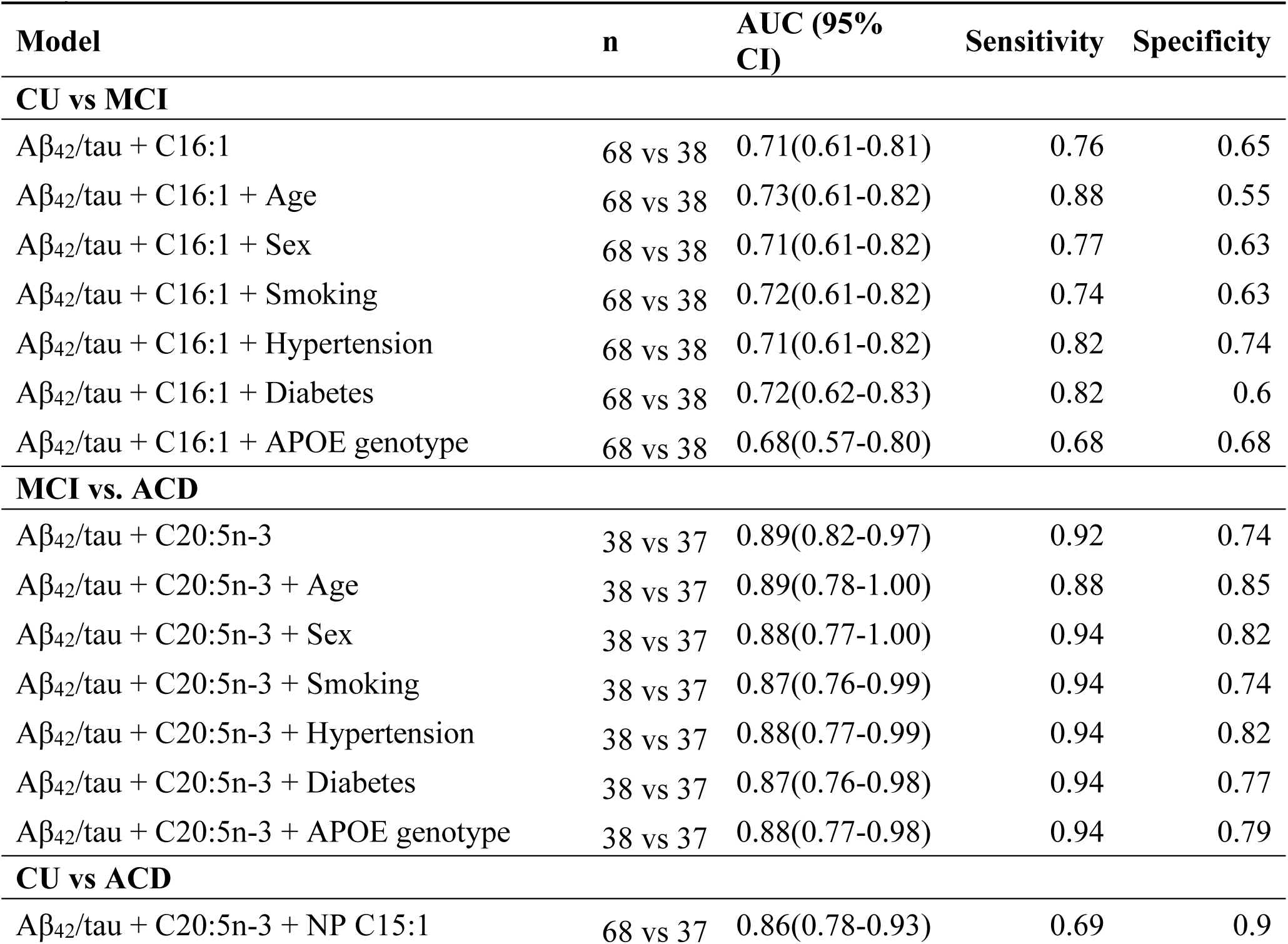
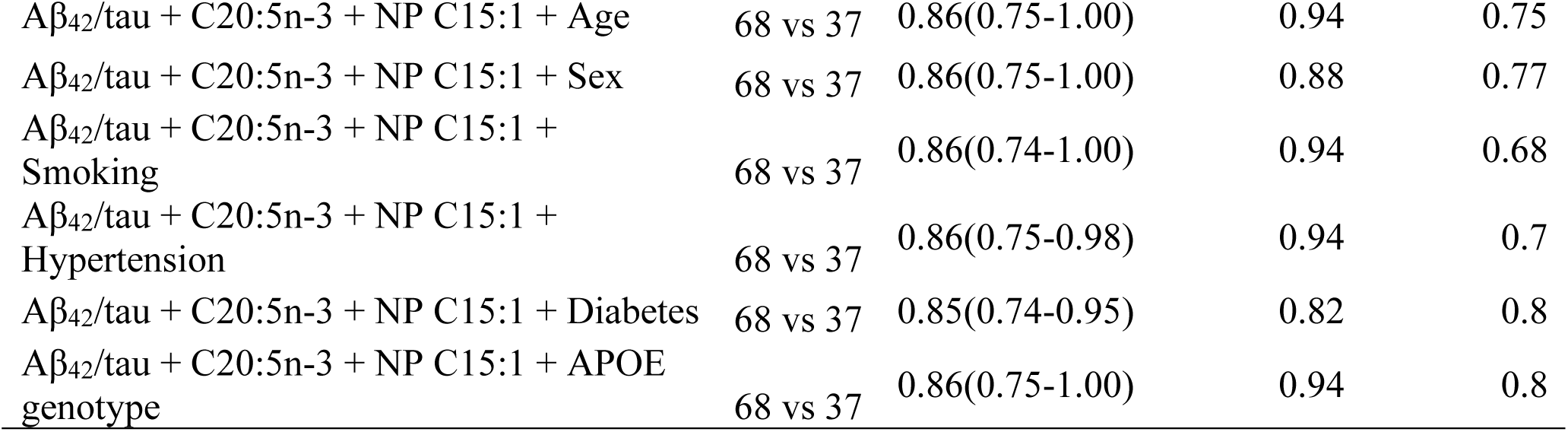
Effect of covariates on the best model for discriminating CU, MCI, and ACD for NP fatty acids.

| Model | n | AUC (95% CI) | Sensitivity | Specificity |
| --- | --- | --- | --- | --- |
| **CU vs MCI** |  |  |  |  |
| A$\beta_{42}$/tau + C16:1 | 68 vs 38 | 0.71(0.61-0.81) | 0.76 | 0.65 |
| A$\beta_{42}$/tau + C16:1 + Age | 68 vs 38 | 0.73(0.61-0.82) | 0.88 | 0.55 |
| A$\beta_{42}$/tau + C16:1 + Sex | 68 vs 38 | 0.71(0.61-0.82) | 0.77 | 0.63 |
| A$\beta_{42}$/tau + C16:1 + Smoking | 68 vs 38 | 0.72(0.61-0.82) | 0.74 | 0.63 |
| A$\beta_{42}$/tau + C16:1 + Hypertension | 68 vs 38 | 0.71(0.61-0.82) | 0.82 | 0.74 |
| A$\beta_{42}$/tau + C16:1 + Diabetes | 68 vs 38 | 0.72(0.62-0.83) | 0.82 | 0.6 |
| A$\beta_{42}$/tau + C16:1 + APOE genotype | 68 vs 38 | 0.68(0.57-0.80) | 0.68 | 0.68 |
| **MCI vs. ACD** |  |  |  |  |
| A$\beta_{42}$/tau + C20:5n-3 | 38 vs 37 | 0.89(0.82-0.97) | 0.92 | 0.74 |
| A$\beta_{42}$/tau + C20:5n-3 + Age | 38 vs 37 | 0.89(0.78-1.00) | 0.88 | 0.85 |
| A$\beta_{42}$/tau + C20:5n-3 + Sex | 38 vs 37 | 0.88(0.77-1.00) | 0.94 | 0.82 |
| A$\beta_{42}$/tau + C20:5n-3 + Smoking | 38 vs 37 | 0.87(0.76-0.99) | 0.94 | 0.74 |
| A$\beta_{42}$/tau + C20:5n-3 + Hypertension | 38 vs 37 | 0.88(0.77-0.99) | 0.94 | 0.82 |
| A$\beta_{42}$/tau + C20:5n-3 + Diabetes | 38 vs 37 | 0.87(0.76-0.98) | 0.94 | 0.77 |
| A$\beta_{42}$/tau + C20:5n-3 + APOE genotype | 38 vs 37 | 0.88(0.77-0.98) | 0.94 | 0.79 |
| **CU vs ACD** |  |  |  |  |
| A$\beta_{42}$/tau + C20:5n-3 + NP C15:1 | 68 vs 37 | 0.86(0.78-0.93) | 0.69 | 0.9 |
| Aβ<sub>42</sub>/tau + C20:5n-3 + NP C15:1 + Age | 68 vs 37 | 0.86(0.75-1.00) | 0.94 | 0.75 |
| Aβ<sub>42</sub>/tau + C20:5n-3 + NP C15:1 + Sex | 68 vs 37 | 0.86(0.75-1.00) | 0.88 | 0.77 |
| Aβ<sub>42</sub>/tau + C20:5n-3 + NP C15:1 +<br>Smoking | 68 vs 37 | 0.86(0.74-1.00) | 0.94 | 0.68 |
| Aβ<sub>42</sub>/tau + C20:5n-3 + NP C15:1 +<br>Hypertension | 68 vs 37 | 0.86(0.75-0.98) | 0.94 | 0.7 |
| Aβ<sub>42</sub>/tau + C20:5n-3 + NP C15:1 + Diabetes | 68 vs 37 | 0.85(0.74-0.95) | 0.82 | 0.8 |
| Aβ<sub>42</sub>/tau + C20:5n-3 + NP C15:1 + APOE<br>genotype | 68 vs 37 | 0.86(0.75-1.00) | 0.94 | 0.8 |

### Summary statistics of the best CSF biomarkers for differential diagnosis of CU, MCI, and ACD

Table 5 summarizes the best-performing CSF fatty acid biomarkers for differential diagnosis of CU, MCI, and ACD. Except MUFA/PUFA, SF C20:1, SF PUFA, NP C14:1, SF C18:3(n-6), and

**Table 5:**
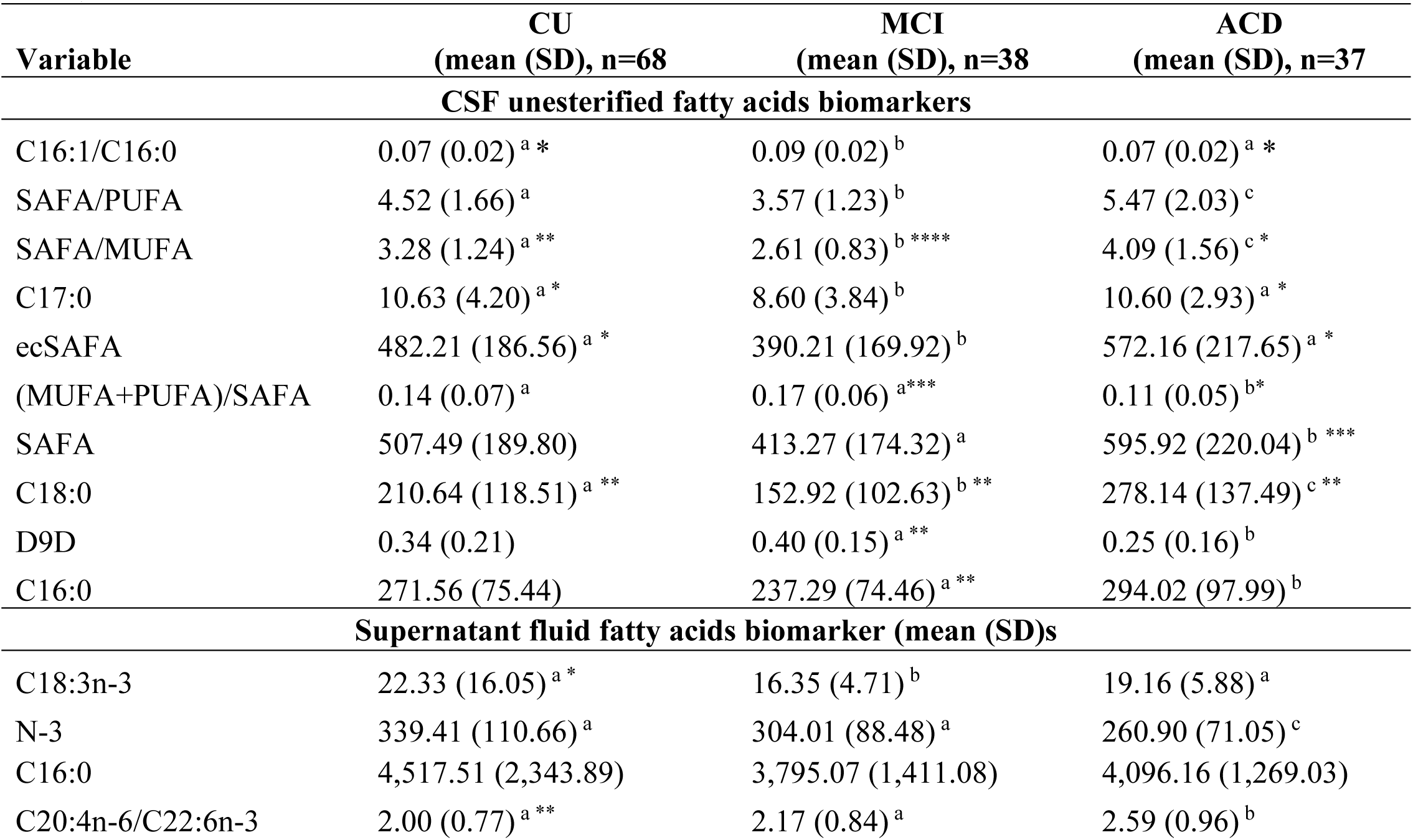
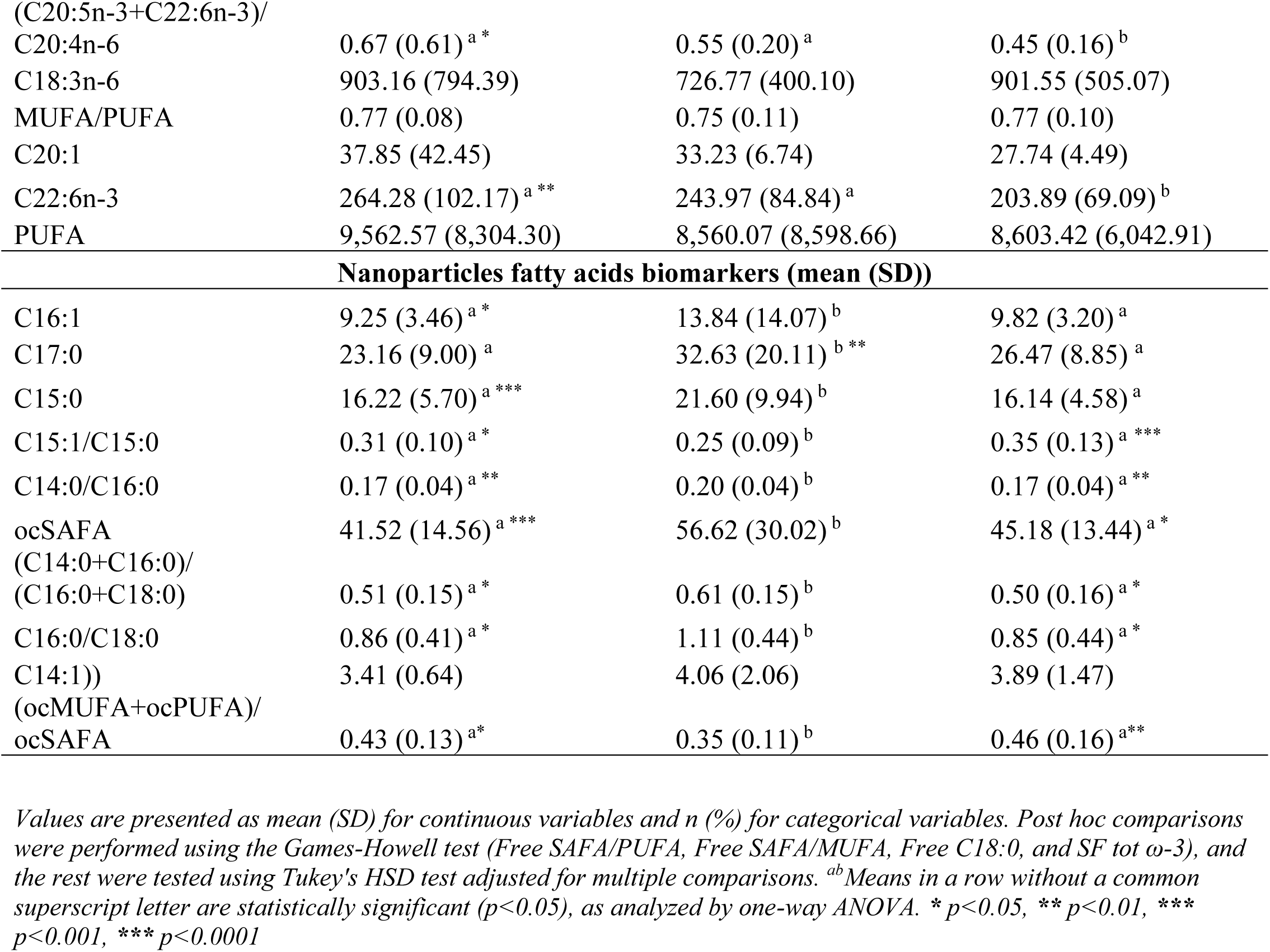
Summary statistics of the best performing CSF fatty acids for the differential diagnosis of CU, MCI, and ACD.

SF C16:0, all the other CSF fractions of fatty acid biomarkers were statistically significantly different between the studied groups (Table 5).

## Discussion

While cerebrospinal fluid (CSF) Aβ_42_ is recognized as an early biomarker of all-cause dementia (ACD), recent studies have identified cognitively normal (CU) individuals to have CSF Aβ_42_ concentrations and Aβ_42_/tau ratios similar to symptomatic ACD patients (Harrington et al., 2013; Sun et al., 2023). Additionally, amyloid deposits have been observed in frontotemporal dementia (FTD) and Lewy Body Dementia (LBD) patients at autopsy (Paterson et al., 2018; Tan et al., 2017), challenging the diagnostic utility of Aβ_42_ and Aβ_42_/tau in ACD. These findings emphasize the need for novel biomarkers to better differentiate between ACD and other dementia subtypes.

Building on the discovery that CSF lipids are distributed between supernatant fluid and brain-derived nanoparticles (Harrington et al., 2009), we hypothesized that fatty acid metabolism in these compartments is altered in early ACD pathology. This study explored the predictive value of CSF unesterified fatty acids, supernatant fluid fatty acids, and nanoparticle fatty acids in the differential diagnosis of CU, mild cognitive impairment (MCI), and ACD.

Using multivariable binary logistic regression and receiver operating characteristic (ROC) curve analysis, our findings demonstrated that CSF fatty acids are clinically relevant biomarkers. Key findings include: (a) Differentiating CU from MCI: Ten biomarkers each from the unesterified, supernatant fluid, and nanoparticle fractions showed superior diagnostic performance compared to Aβ_42_, tau, and Aβ_42_/tau; (b) Differentiating MCI from ACD and CU from ACD: Aβ_42_, tau, and Aβ_42_/tau outperformed CSF fatty acids, though the differences in diagnostic accuracy were not statistically significant; (c) Combination biomarkers: Integrating CSF fatty acids with the Aβ_42_/tau ratio improved diagnostic accuracy; (d) Covariate influence: Demographic and clinical variables such as age, APOE genotype, sex, and comorbidities had no significant impact on the diagnostic utility of fatty acids.

These findings suggest that changes in CSF fatty acid composition precede canonical ACD biomarkers in detecting the transition from CU to MCI. We found CSF-fraction-specific differences in fatty acids that classified CU and MCI. Specifically, for the unesterified fatty acid fraction, the desaturase index (DI), such as C16:1/C16:0, the saturation index (SAFA/PUFA), and levels of saturated fatty acids (C16:0, C17:0, C18:0, even-chain SAFA) showed strong discriminatory power. In contrast, omega-3 fatty acids (C18:3n-3, n-3, C20:3n-3, PUFA), the inflammatory index (AA/DHA), the anti-inflammatory index (EPA+DHA)/AA), a SAFA (C16:0), and a MUFA (C20:1) were top classifiers of CU versus MCI in the supernatant fluid. For the nanoparticulate fraction, desaturase index of even-chain (C16:1/C16:0) and odd-chain fatty acids (C15:1/C5:0, ocDI), odd-chain SAFA, and elongase indices were the top classifiers of CU versus MCI (Fig. 4A). These biomarkers may provide unique insights into lipid metabolic changes linked to early ACD pathology since Aβ_42_ and tau became the major classifiers only when comparing CU versus AD and MCI versus AD (Fig. 4A).

**Fig. 4.**
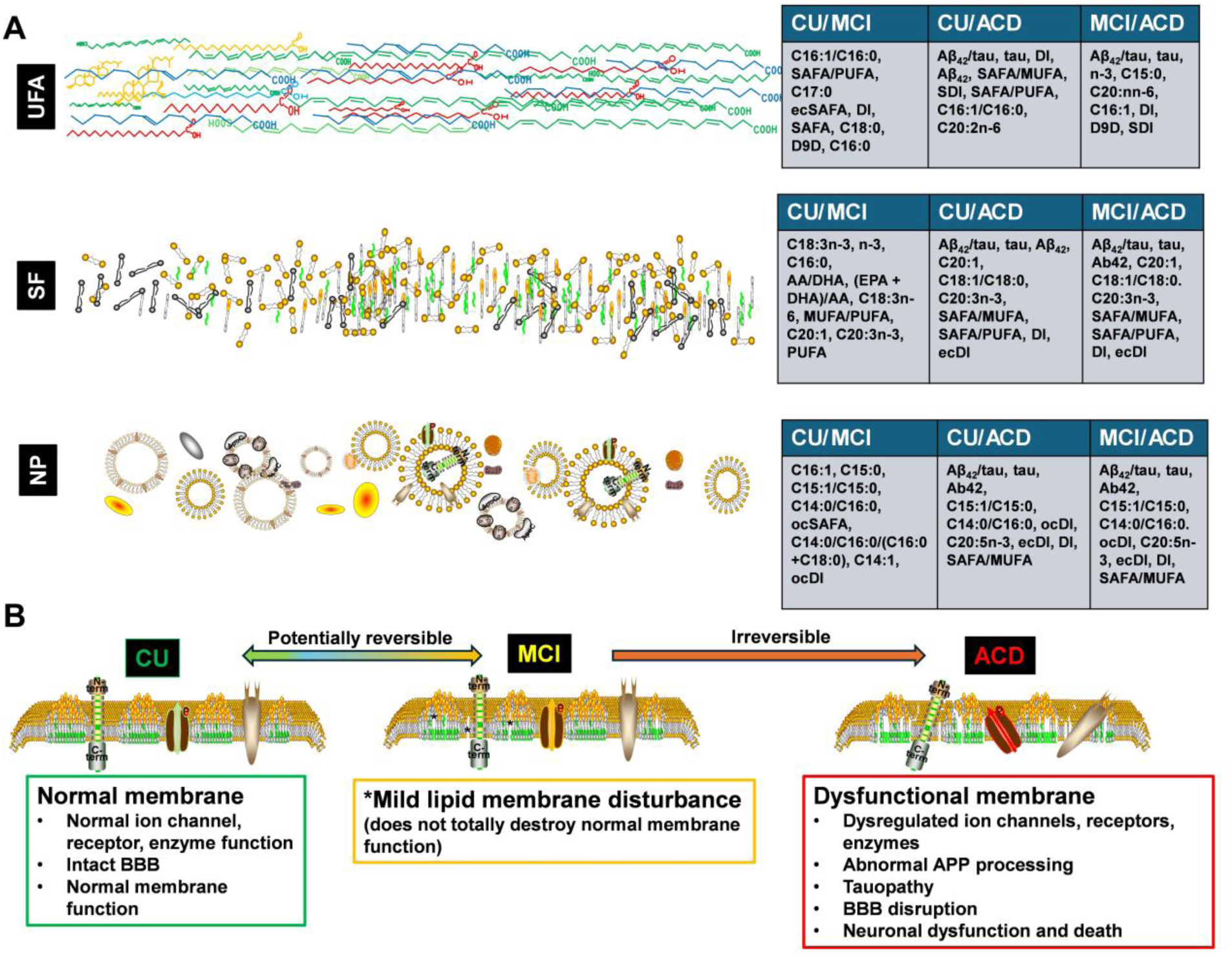
The top classifiers of CU versus MCI, CU versus ACD, and MCI versus ACD in CSF membrane fractions (UFA, unesterified fatty acids; SF supernatant fluid; NP, nanoparticles). For CU versus MCI, the top classifiers are lipids while Aβ_42_/tau are top classifiers in CU versus ACD and MCI versus ACD (Fig. 4A). The changes in membrane lipids my result in dysfunctional properties associated with ACD pathology (Fig. 4B).

We hypothesize that the early alteration in lipid metabolism initiates a pathological process that may eventually result in abnormal amyloid precursor protein processing, tauopathy, disrupted blood-brain barrier, dysregulated ion channels and receptors, and eventually neuronal and axonal degeneration (Fig. 4B).

Our results align with previous studies emphasizing the value of multivariable biomarker panels. For example, Thijssen et al., (2022) highlighted the diagnostic potential of combining Aβ_42_/40, p-tau181, NfL, and GFAP to distinguish ACD from FTD and DLB. Similarly, Krishna et al., (2024) demonstrated improved diagnostic accuracy using NfL/Aβ_42_, t-tau, and p-tau181. Consistent with this approach, our study revealed that incorporating CSF fatty acids with Aβ_42_/tau significantly enhances diagnostic performance, particularly in differentiating CU from MCI. For example, the AUC of Aβ_42_/tau increased from 0.55 to 0.69-0.71 with unesterified fatty acids and from 0.55 to 0.67 with supernatant fatty acids.

Our findings underscore the importance of lipid metabolism in early ACD pathology. Notably, reductions in supernatant DHA, alpha-linoleic acid, total omega-3 fatty acids, and nanoparticle EPA were observed in ACD compared to CU. These differences may reflect lipid peroxidation, reacylation, phospholipase A_2_ hydrolysis, and impaired uptake into the brain (Dakterzada et al., 2024; Yin, 2023). Prior research suggests that omega-3 fatty acids such as DHA and EPA have neuroprotective properties, including inhibiting Aβ synthesis, enhancing its degradation, and modulating β- and γ-secretase activity (Grimm et al., 2011). In AD mouse models, palmitic acid levels were elevated, consistent with overexpression of APP, PSEN1, and acetyl-CoA carboxylase (Ates et al., 2020). We also noticed that odd-chain fatty acids (C15:0, C17:0) were good early biomarkers of ACD. Several studies have identified ocFA as essential for several physiological functions (Chen et al., 2025; Pfeuffer & Jaudszus, 2016; Yang et al., 2025). These ocFAs may be derived from microbial metabolism (Qin et al., 2023; Zhang et al., 2020) and thus highlight the important role of the microbiome in cognitive function.

### Strengths and Limitations

This study provides novel insights into the early lipid metabolic changes in ACD pathology, highlighting the diagnostic potential of CSF fatty acids. It is among the few studies to explore CSF fatty acids as biomarkers across the CU-MCI-ACD continuum. Additionally, the lack of considerable influence from covariates demonstrates the robustness of these biomarkers in the tightly regulated brain environment. However, the study’s moderate sample size limits generalizability, and the cross-sectional design precludes causal inferences. Moreover, plasma fatty acid concentrations and neuroimaging data were unavailable, preventing a direct comparison of CSF and plasma biomarkers. The absence of CSF Aβ40, p-tau217, and p-tau218 data limited the scope of biomarker comparisons. Future studies should validate these findings in larger, longitudinal cohorts and evaluate the utility of CSF fatty acids in combination with advanced neuroimaging and plasma markers.

## Conclusion

Fatty acids and their desaturation indices in CSF fractions (unesterified fatty acids, supernatant fluid fatty acids, nanoparticle) have emerged as superior classifiers for distinguishing cognitively unimpaired (CU) individuals from those with mild cognitive impairment (MCI). However, Aβ, and tau remain the most effective biomarkers for differentiating between MCI and all-cause Alzheimer’s disease (ACD), as well as CU and ACD. Notably, their discriminative power does not show statistically significant differences when compared to the fatty acid biomarkers identified in CSF fractions.

These findings collectively suggest that disruptions in fatty acid metabolism may precede amyloid and tau abnormalities in the initial stages of ACD pathology. Consequently, early metabolic alterations in fatty acids may serve as more sensitive indicators of early cognitive impairment. This highlights the potential for therapeutic strategies targeting fatty acid metabolism to mitigate cognitive decline in aging populations.

Furthermore, this study underscores the value of integrating multivariable analyses of canonical ACD biomarkers with CSF fatty acid profiles to enhance the diagnostic accuracy of ACD across its clinical spectrum. Future research should prioritize controlled longitudinal studies to identify novel CSF fatty acid biomarkers, elucidate their role in cognitive function, and clarify their contributions to ACD pathophysiology. Such studies could pave the way for a deeper understanding of the metabolic mechanisms underlying ACD and the development of early interventions.

## List of Abbreviations

ACD: All-cause dementia
MCI: Mild Cognitive Impairment
CU: Cognitively Unimpaired
CSF: Cerebrospinal fluid
Aβ42: Amyloid beta 42
FTD: Frontotemporal dementia
DLB: Lewy body dementia
NfL: Neurofilament light chain
GFAP: Glial fibrillary acidic protein
p-tau181: Phosphorylated tau 181
BBB: Blood brain barrier
APP: Amyloid precursor protein
LysoPC/PC: lysophosphatidylcholine/phosphatidylcholine
t-tau: Total tau
SAFA: Saturated fatty acid
ocSAFA: Odd chain saturated fatty acid
EcSAFA: Even chain saturated fatty acid
MUFA: Monounsaturated fatty acid
ocMUFA: Odd chain monounsaturated fatty acid
ecMUFA: Even chain monosaturated fatty acid
PUFA: Polyunsaturated fatty acid
ocPUFA: Odd chain polyunsaturated fatty acid
EcPUFA: Even chain polyunsaturated fatty acid
D9D: Delta-9 Desaturase
DI: Desaturase Index
OcDI: Odd-chain Desaturase Index
EcDI: Even-chain Desaturase Index
SDI: Stearoyl-CoA Desaturation Index
ROC: Receiver Operating Characteristics
AUC: Area Under the Curve

## Acknowledgment

The authors thank Huntington Medical Research Institutes for providing the data. They are grateful to the African Initiative on Bioinformatics Online Training on Neurodegenerative Diseases (AI-BOND) for the training and support offered to Jacob Apibilla Ayembilla.

## Authors’ contribution

JAA, YNW, and RB analyzed the data and wrote the draft manuscript, BAA, ABSND, MS, XJ, AKN, SS, JJH, BF, AF conceptualized the project, AF provided the data. All authors read, reviewed, and approved the final manuscript for submission.

## Ethics approval and consent to participate

Ethical approval for the study was previously obtained by the Huntington Medical Research Institute. Since this study is a secondary analysis of their data, permission was obtained to use the data.

## Consent for publication

Not applicable

## Availability of data and materials

The datasets used and/or analyzed during the current study are available from the corresponding author on reasonable request.

## References

Ates, G., Goldberg, J., Currais, A., & Maher, P. (2020). CMS121, a fatty acid synthase inhibitor, protects against excess lipid peroxidation and inflammation and alleviates cognitive loss in a transgenic mouse model of Alzheimer’s disease. Redox Biology, 36, 101648. 10.1016/j.redox.2020.101648

Bousiges, O., & Blanc, F. (2022). Biomarkers of Dementia with Lewy Bodies: Differential Diagnostic with Alzheimer’s Disease. International Journal of Molecular Sciences, 23(12). 10.3390/ijms23126371

Brand, A. L., Lawler, P. E., Bollinger, J. G., Li, Y., Schindler, S. E., Li, M., Lopez, S., Ovod, V., Nakamura, A., Shaw, L. M., Zetterberg, H., Hansson, O., & Bateman, R. J. (2022). The performance of plasma amyloid beta measurements in identifying amyloid plaques in Alzheimer’s disease: a literature review. Alzheimer’s Research and Therapy, 14(1), 1–15. 10.1186/s13195-022-01117-1

Cardoso, C., Afonso, C., & Bandarra, N. M. (2016). Dietary DHA and health: Cognitive function ageing. Nutrition Research Reviews, 29(2), 281–294. 10.1017/S0954422416000184

Castor, K. J., Shenoi, S., Edminster, S. P., Tran, T., King, K. S., Chui, H., Pogoda, J. M., Fonteh, A. N., & Harrington, M. G. (2020). Urine dicarboxylic acids change in presymptomatic Alzheimer’s disease and reflect loss of energy capacity and hippocampal volume. PLoS ONE, 15(4). 10.1371/journal.pone.0231765

Cederholm, T., Salem, N., & Palmblad, J. (2013). Ω-3 Fatty Acids in the Prevention of Cognitive Decline in Humans. Advances in Nutrition, 4(6), 672–676. 10.3945/an.113.004556

Chen, M. K., Mecca, A. P., Naganawa, M., Finnema, S. J., Toyonaga, T., Lin, S. F., Najafzadeh, S., Ropchan, J., Lu, Y., McDonald, J. W., Michalak, H. R., Nabulsi, N. B., Arnsten, A. F. T., Huang, Y., Carson, R. E., & Van Dyck, C. H. (2018). Assessing Synaptic Density in Alzheimer Disease with Synaptic Vesicle Glycoprotein 2A Positron Emission Tomographic Imaging. JAMA Neurology, 75(10), 1215–1224. 10.1001/jamaneurol.2018.1836

Cutuli, D., Landolfo, E., Decandia, D., Nobili, A., Viscomi, M. T., La Barbera, L., Sacchetti, S., De Bartolo, P., Curci, A., D’Amelio, M., Farioli-Vecchioli, S., & Petrosini, L. (2022). Correction: Cutuli et al. Neuroprotective Role of Dietary Supplementation with Omega-3 Fatty Acids in the Presence of Basal Forebrain Cholinergic Neurons Degeneration in Aged Mice (Int. J. Mol. Sci. 2020, 21, 1741). International Journal of Molecular Sciences, 23(13). 10.3390/ijms23136916

Dakterzada, F., Jové, M., Huerto, R., Carnes, A., Sol, J., Pamplona, R., & Piñol-Ripoll, G. (2024). Cerebrospinal fluid neutral lipids predict progression from mild cognitive impairment to Alzheimer’s disease. GeroScience, 46(1), 683–696. 10.1007/s11357-023-00989-x

Davidson, C. G., Woodford, S. J., Mathur, S., Valle, D. B., Kioutchoukova, I., Mahmood, A., & Lucke-wold, B. (2023). Exploration of Neuroscience Investigation into the vascular contributors to dementia and the associated treatments. 224–237. 10.37349/en.2023.00023

Davis, A., Mendoza, W., Leach, D., & Marques, O. (2022). Predicting Alzheimer’s disease with multi-omic data: A systematic review. MedRxiv. 10.1101/2022.11.25.22282770

Di Miceli, M., Martinat, M., Rossitto, M., Aubert, A., Alashmali, S., Bosch-bouju, C., Fioramonti, X., Joffre, C., Bazinet, R. P., & Layé, S. (2022). Dietary Long-Chain n-3 Polyunsaturated Fatty Acid Supplementation Alters Electrophysiological Properties in the Nucleus Accumbens and Emotional Behavior in Naïve and Chronically Stressed Mice. International Journal of Molecular Sciences, 23(12). 10.3390/ijms23126650

Dong, X., Nao, J., Shi, J., & Zheng, D. (2019). Predictive Value of Routine Peripheral Blood Biomarkers in Alzheimer’s Disease. Frontiers in Aging Neuroscience, 11(December), 1–9. 10.3389/fnagi.2019.00332

Dubois, B., Epelbaum, S., Nyasse, F., Bakardjian, H., Gagliardi, G., Uspenskaya, O., Houot, M., Lista, S., Cacciamani, F., Potier, M. C., Bertrand, A., Lamari, F., Benali, H., Mangin, J. F., Colliot, O., Genthon, R., Habert, M. O., Hampel, H., Audrain, C., … Younsi, N. (2018). Cognitive and neuroimaging features and brain β-amyloidosis in individuals at risk of Alzheimer’s disease (INSIGHT-preAD): a longitudinal observational study. The Lancet Neurology, 17(4), 335–346. 10.1016/S1474-4422(18)30029-2

Dubois, B., von Arnim, C. A. F., Burnie, N., Bozeat, S., & Cummings, J. (2023). Biomarkers in Alzheimer’s disease: role in early and differential diagnosis and recognition of atypical variants. Alzheimer’s Research and Therapy, 15(1), 1–13. 10.1186/s13195-023-01314-6

Fonteh, A. N., Cipolla, M., Chiang, A. J., Edminster, S. P., Arakaki, X., & Harrington, M. G. (2020). Polyunsaturated Fatty Acid Composition of Cerebrospinal Fluid Fractions Shows Their Contribution to Cognitive Resilience of a Pre-symptomatic Alzheimer’s Disease Cohort. Frontiers in Physiology, 11(February), 1–14. 10.3389/fphys.2020.00083

Fonteh, A. N., Cipolla, M., Chiang, J., Arakaki, X., & Harrington, M. G. (2014). Human cerebrospinal fluid fatty acid levels differ between supernatant fluid and brain-derived nanoparticle fractions, and are altered in Alzheimer’s disease. PLoS ONE, 9(6), 1–14. 10.1371/journal.pone.0100519

Grimm, M. O. W., Kuchenbecker, J., Grosgen, S., Burg, V. K., Hundsdorfer, B., Rothhaar, T. L., Friess, P., De Wilde, M. C., Broersen, L. M., Penke, B., Peter, M., Vígh, L., Grimm, H. S., & Hartmann, T. (2011). Docosahexaenoic acid reduces amyloid β production via multiple pleiotropic mechanisms. Journal of Biological Chemistry, 286(16), 14028–14039. 10.1074/jbc.M110.182329

Harrington, M. G., Chiang, J., Pogoda, J. M., Gomez, M., Thomas, K., Marion, S. D. B., Miller, K. J., Siddarth, P., Yi, X., Zhou, F., Lee, S., Arakaki, X., Cowan, R. P., Tran, T., Charleswell, C., Ross, B. D., & Fonteh, A. N. (2013). Executive function changes before memory in preclinical Alzheimer’s pathology: A prospective, cross-sectional, case control study. PLoS ONE, 8(11). 10.1371/journal.pone.0079378

Harrington, M. G., Edminster, S. P., Buennagel, D. P., Chiang, J. P., Sweeney, M. D., Chui, H. C., Zlokovic, B. V., & Fonteh, A. N. (2019). P1-165: Four-Year Longitudinal Study of Cognitively Healthy Individuals: Csf Amyloid/Tau Levels and Nanoparticle Membranes Identify High Risk for Alzheimer’S Disease. Alzheimer’s & Dementia, 15(7S_Part_5), P299. 10.1016/j.jalz.2019.06.720

Harrington, M. G., Fonteh, A. N., Oborina, E., Liao, P., Cowan, R. P., McComb, G., Chavez, J. N., Rush, J., Biringer, R. G., & Hühmer, A. F. (2009). The morphology and biochemistry of nanostructures provide evidence for synthesis and signaling functions in human cerebrospinal fluid. Cerebrospinal Fluid Research, 6(1). 10.1186/1743-8454-6-10

Iaccarino, L., Burnham, S. C., Dell’Agnello, G., Dowsett, S. A., & Epelbaum, S. (2023). Diagnostic Biomarkers of Amyloid and Tau Pathology in Alzheimer’s Disease: An Overview of Tests for Clinical Practice in the United States and Europe. Journal of Prevention of Alzheimer’s Disease, 10(3), 426–442. 10.14283/jpad.2023.43

Janelidze, S., Mattsson, N., Palmqvist, S., Smith, R., Beach, T. G., Serrano, G. E., Chai, X., Proctor, N. K., Eichenlaub, U., Zetterberg, H., Blennow, K., Reiman, E. M., Stomrud, E., Dage, J. L., & Hansson, O. (2020). Plasma P-tau181 in Alzheimer’s disease: relationship to other biomarkers, differential diagnosis, neuropathology and longitudinal progression to Alzheimer’s dementia. Nature Medicine, 26(3), 379–386. 10.1038/s41591-020-0755-1

Kashyap, G., Bapat, D., Das, D., Gowaikar, R., Amritkar, R. E., Rangarajan, G., Ravindranath, V., & Ambika, G. (2019). Synapse loss and progress of Alzheimer’s disease -A network model. Scientific Reports, 9(1). 10.1038/s41598-019-43076-y

Kosicek, M., Zetterberg, H., Andreasen, N., Peter-Katalinic, J., & Hecimovic, S. (2012). Elevated cerebrospinal fluid sphingomyelin levels in prodromal Alzheimer’s disease. Neuroscience Letters, 516(2), 302–305. 10.1016/j.neulet.2012.04.019

Krishna, G., Thangaraju Sivakumar, P., Dahale, A. B., & Subramanian, S. (2024). Potential Utility of Plasma Biomarker Panels in Differential Diagnosis of Alzheimer’s Disease. Journal of Alzheimer’s Disease Reports, 8(1), 1–7. 10.3233/ADR-230156

Kuhn, M. (2008). Building predictive models in R using the caret package. Journal of Statistical Software, 28(5), 1–26. 10.18637/jss.v028.i05

Liu, Y., Wei, M., Yue, K., Hu, M., Li, S., Men, L., Pi, Z., Liu, Z., & Liu, Z. (2018). Study on Urine Metabolic Profile of Aβ25–35-Induced Alzheimer’s Disease Using UHPLC-Q-TOF-MS. Neuroscience, 394, 30–43. 10.1016/j.neuroscience.2018.10.001

LoBue, C., Munro Cullum, C., Didehbani, N., Yeatman, K., Jones, B., Kraut, M. A., & Hart, J. (2018). Neurodegenerative dementias after traumatic brain injury. Journal of Neuropsychiatry and Clinical Neurosciences, 30(1), 7–13. 10.1176/appi.neuropsych.17070145

Oksman, M., Iivonen, H., Hogyes, E., Amtul, Z., Penke, B., Leenders, I., Broersen, L., Lütjohann, D., Hartmann, T., & Tanila, H. (2006). Impact of different saturated fatty acid, polyunsaturated fatty acid and cholesterol containing diets on beta-amyloid accumulation in APP/PS1 transgenic mice. Neurobiology of Disease, 23(3), 563–572. 10.1016/j.nbd.2006.04.013

Paterson, R. W., Slattery, C. F., Poole, T., Nicholas, J. M., Magdalinou, N. K., Toombs, J., Chapman, M. D., Lunn, M. P., Heslegrave, A. J., Foiani, M. S., Weston, P. S. J., Keshavan, A., Rohrer, J. D., Rossor, M. N., Warren, J. D., Mummery, C. J., Blennow, K., Fox, N. C., Zetterberg, H., & Schott, J. M. (2018). Cerebrospinal fluid in the differential diagnosis of Alzheimer’s disease: Clinical utility of an extended panel of biomarkers in a specialist cognitive clinic. Alzheimer’s Research and Therapy, 10(1), 1–11. 10.1186/s13195-018-0361-3

R Core Team. (2021). R: A language and environment for statistical computing. R foundation for statistical computing. R Foundation for Statistical Computing, 2021. https://www.r-project.org/

Richter, N., Nellessen, N., Dronse, J., Dillen, K., Jacobs, H. I. L., Langen, K. J., Dietlein, M., Kracht, L., Neumaier, B., Fink, G. R., Kukolja, J., & Onur, O. A. (2019). Spatial distributions of cholinergic impairment and neuronal hypometabolism differ in MCI due to AD. NeuroImage: Clinical, 24. 10.1016/j.nicl.2019.101978

Sheng, M., Sabatini, B. L., & Südhof, T. C. (2012). Synapses and Alzheimer’s disease. Cold Spring Harbor Perspectives in Biology, 4(5), 10. 10.1101/cshperspect.a005777

Sun, Y., Zhao, Y., Hu, K., Wang, M., Liu, Y., & Liu, B. (2023). Distinct spatiotemporal subtypes of amyloid deposition are associated with diverging disease profiles in cognitively normal and mild cognitive impairment individuals. Translational Psychiatry, 13(1). 10.1038/s41398-023-02328-2

Tan, R. H., Kril, J. J., Yang, Y., Tom, N., Hodges, J. R., Villemagne, V. L., Rowe, C. C., Leyton, C. E., Kwok, J. B. J., Ittner, L. M., & Halliday, G. M. (2017). Assessment of amyloid β in pathologically confirmed frontotemporal dementia syndromes. *Alzheimer’s and Dementia: Diagnosis*, Assessment and Disease Monitoring, 9, 10–20. 10.1016/j.dadm.2017.05.005

Tao, Q. Q., Lin, R. R., & Wu, Z. Y. (2023). Early Diagnosis of Alzheimer’s Disease: Moving Toward a Blood-Based Biomarkers Era. Clinical Interventions in Aging, 18(February), 353–358. 10.2147/CIA.S394821

Thijssen, E. H., Verberk, I. M. W., Kindermans, J., Abramian, A., Vanbrabant, J., Ball, A. J., Pijnenburg, Y., Lemstra, A. W., van der Flier, W. M., Stoops, E., Hirtz, C., & Teunissen, C. E. (2022). Differential diagnostic performance of a panel of plasma biomarkers for different types of dementia. *Alzheimer’s and Dementia: Diagnosis*, Assessment and Disease Monitoring, 14(1), 1–11. 10.1002/dad2.12285

Tong, B., Ba, Y., Li, Z., Yang, C., Su, K., Qi, H., Zhang, D., Liu, X., Wu, Y., Chen, Y., Ling, J., Zhang, J., Yin, X., & Yu, P. (2024). Targeting dysregulated lipid metabolism for the treatment of Alzheimer’s disease and Parkinson’s disease: Current advancements and future prospects. Neurobiology of Disease, 196(January), 106505. 10.1016/j.nbd.2024.106505

Turck, N., Vutskits, L., Sanchez-Pena, P., Robin, X., Hainard, A., Gex-Fabry, M., Fouda, C., Bassem, H., Mueller, M., Lisacek, F., Puybasset, L., & Sanchez, J.-C. (2011). pROC: an open-source package for R and S+ to analyze and compare ROC curves. BMC Bioinformatics, 8, 12–77. http://link.springer.com/10.1007/s00134-009-1641-y

Walter, A., Korth, U., Hilgert, M., Hartmann, J., Weichel, O., Hilgert, M., Fassbender, K., Schmitt, A., & Klein, J. (2004). Glycerophosphocholine is elevated in cerebrospinal fluid of Alzheimer patients. Neurobiology of Aging, 25(10), 1299–1303. 10.1016/j.neurobiolaging.2004.02.016

Whiley, L., Sen, A., Heaton, J., Proitsi, P., García-Gómez, D., Leung, R., Smith, N., Thambisetty, M., Kloszewska, I., Mecocci, P., Soininen, H., Tsolaki, M., Vellas, B., Lovestone, S., & Legido-Quigley, C. (2014). Evidence of altered phosphatidylcholine metabolism in Alzheimer’s disease. Neurobiology of Aging, 35(2), 271–278. 10.1016/j.neurobiolaging.2013.08.001

Yin, F. (2023). Lipid metabolism and Alzheimer’s disease: clinical evidence, mechanistic link and therapeutic promise. 290(6), 1420–1453. 10.1111/febs.16344.Lipid

## REFERENCES

Chen, T., Luo, J., Li, S., Li, X., Wang, W., Lu, W.,…Xu, X. (2025). Associations between serum pentadecanoic acid (C15:0) and heptadecanoic acid (C17:0) levels and hypertension: a cross-sectional analysis of NHANES data. Lipids Health Dis, 24(1), 219. 10.1186/s12944-025-02640-4

Pfeuffer, M., & Jaudszus, A. (2016). Pentadecanoic and Heptadecanoic Acids: Multifaceted Odd-Chain Fatty Acids. Adv Nutr, 7(4), 730–734. 10.3945/an.115.011387

Qin, N., Li, L., Wang, Z., & Shi, S. (2023). Microbial production of odd-chain fatty acids. Biotechnol Bioeng, 120(4), 917–931. 10.1002/bit.28308

Yang, Y., Fu, Y., & Wu, C. (2025). Gut microbe-derived pentadecanoic acid could represent a novel health-promoter. Food Funct, 16(12), 4636–4653. 10.1039/d5fo01278c

Zhang, L. S., Liang, S., Zong, M. H., Yang, J. G., & Lou, W. Y. (2020). Microbial synthesis of functional odd-chain fatty acids: a review. World J Microbiol Biotechnol, 36(3), 35. 10.1007/s11274-020-02814-5

