## Supplementary Data for "Early Metabolic Alterations in Cerebrospinal Fluid Fatty Acid Profiles Linked to Cognitive Decline and All-Cause Dementia"

**Table S1: Best performing CSF unesterified fatty acids for differential diagnosis of CU, MCI, and ACD based on ROC curve analyses.**

| Variable | n | AUC (95% CI) | Sensitivity | Specificity |
|---|---|---|---|---|
| **CU versus MCI** | | | | |
| C16:1/FA16:0 | 68 vs 38 | 0.70(0.60-0.81) | 0.68 | 0.75 |
| SAFA/PUFA | 68 vs 38 | 0.69(0.59-0.80) | 0.55 | 0.81 |
| SAFA/MUFA | 68 vs 38 | 0.69(0.59-0.80) | 0.66 | 0.72 |
| C17:0 | 68 vs 38 | 0.68(0.58-0.79) | 0.58 | 0.78 |
| DI | 68 vs 38 | 0.67(0.56-0.78) | 0.74 | 0.6 |
| ecSAFA | 68 vs 38 | 0.67(0.56-0.77) | 0.45 | 0.88 |
| SAFA | 68 vs 38 | 0.67(0.56-0.78) | 0.45 | 0.87 |
| C18:0 | 68 vs 38 | 0.67(0.56-0.78) | 0.71 | 0.59 |
| D9D | 68 vs 38 | 0.67(0.56-0.77) | 0.76 | 0.57 |
| C16:0 | 68 vs 38 | 0.66(0.55-0.77) | 0.39 | 0.9 |
| **MCI versus ACD** | | | | |
| Aβ42/tau | 38 vs 37 | 0.85(0.75-0.94) | 0.89 | 0.76 |
| Tau | 38 vs 37 | 0.82(0.72-0.92) | 0.73 | 0.84 |
| D9D | 38 vs 37 | 0.79(0.68-0.90) | 0.73 | 0.76 |
| Aβ42 | 38 vs 37 | 0.79(0.68-0.89) | 0.78 | 0.82 |
| DI | 38 vs 37 | 0.79(0.68-0.89) | 0.78 | 0.71 |
| SAFA/MUFA | 38 vs 37 | 0.78(0.67-0.89) | 0.65 | 0.87 |
| SDI | 38 vs 37 | 0.78(0.67-0.89) | 0.73 | 0.76 |
| SAFA/PUFA | 38 vs 37 | 0.77(0.66-0.88) | 0.68 | 0.84 |
| C16:1/C16:0 | 38 vs 37 | 0.77(0.66-0.89) | 0.76 | 0.79 |
| C20:2(n-6) | 38 vs 37 | 0.77(0.66-0.88) | 0.73 | 0.76 |
| **CU versus ACD** | | | | |
| Aβ42/tau | 68 vs 37 | 0.85(0.77-0.93) | 0.87 | 0.75 |
| Tau | 68 vs 37 | 0.80(0.72-0.89) | 0.84 | 0.71 |
| Aβ42 | 68 vs 37 | 0.76(0.67-0.86) | 0.76 | 0.74 |
| N-3 | 68 vs 37 | 0.69(0.58-0.80) | 0.57 | 0.79 |
| C15:0 | 68 vs 37 | 0.68(0.57-0.79) | 0.41 | 0.88 |
| C20:2(n-6) | 68 vs 37 | 0.68(0.56-0.79) | 0.51 | 0.85 |
| C16:1 | 68 vs 37 | 0.68(0.57-0.78) | 0.57 | 0.75 |
| DI | 68 vs 37 | 0.67(0.55-0.78) | 0.6 | 0.78 |

| | | | | |
|---|---|---|---|---|
| D9D | 68 vs 37 | 0.66(0.55-0.78) | 0.51 | 0.85 |
| SDI | 68 vs 37 | 0.66(0.54-0.78) | 0.51 | 0.87 |

**Table S2: DeLong test for statistically significant difference between the AUC, sensitivity, and specificity of CSF unesterified fatty acids and Aβ42, tau, and Aβ42/tau**

| Predictor1 | Predictor2 | p-value |
|---|---|---|
| **CU vs MCI** | | |
| $A\beta_{42}$ | C16:1/C16:0 | 0.034 |
| $A\beta_{42}$ | SAFA/MUFA | 0.042 |
| $A\beta_{42}$ | SAFA/PUFA | 0.038 |
| Tau | C16:1/C16:0 | 0.034 |
| Tau | SAFA/MUFA | 0.036 |
| Tau | SAFA/PUFA | 0.039 |
| $A\beta_{42}$/tau | C16:1/C16:0 | >0.05 |
| $A\beta_{42}$/tau | SAFA/MUFA | >0.05 |
| $A\beta_{42}$/tau | SAFA/PUFA | >0.05 |
| $A\beta_{42}$/tau | DI | >0.05 |
| Tau | DI | >0.05 |
| $A\beta_{42}$ | DI | >0.05 |
| $A\beta_{42}$/tau | C17:0 | >0.05 |
| Tau | C17:0 | >0.05 |
| $A\beta_{42}$ | C17:0 | >0.05 |
| $A\beta_{42}$/tau | ecSAFA | >0.05 |
| Tau | ecSAFA | >0.05 |
| $A\beta_{42}$ | ecSAFA | >0.05 |
| $A\beta_{42}$/tau | SAFA | >0.05 |
| Tau | SAFA | >0.05 |
| $A\beta_{42}$ | SAFA | >0.05 |
| $A\beta_{42}$/tau | C18:0 | >0.05 |
| Tau | C18:0 | >0.05 |
| $A\beta_{42}$ | C18:0 | >0.05 |
| $A\beta_{42}$/tau | D9D | >0.05 |
| Tau | D9D | >0.05 |
| $A\beta_{42}$ | D9D | >0.05 |
| $A\beta_{42}$/tau | C16:0 | >0.05 |
| Tau | C16:0 | >0.05 |
| $A\beta_{42}$ | C16:0 | >0.05 |
| **MCI vs ACD** | | |
| $A\beta_{42}$/tau | D9D | >0.05 |

| | | |
|---|---|---|
| Tau | D9D | >0.05 |
| $A\beta_{42}$ | D9D | >0.05 |
| $A\beta_{42}$/tau | DI | >0.05 |
| Tau | DI | >0.05 |
| $A\beta_{42}$ | DI | >0.05 |
| $A\beta_{42}$/tau | SAFA/MUFA | >0.05 |
| Tau | SAFA/MUFA | >0.05 |
| $A\beta_{42}$ | SAFA/MUFA | >0.05 |
| $A\beta_{42}$/tau | SDI | >0.05 |
| Tau | SDI | >0.05 |
| $A\beta_{42}$ | SDI | >0.05 |
| $A\beta_{42}$/tau | SAFA/PUFA | >0.05 |
| Tau | SAFA/PUFA | >0.05 |
| $A\beta_{42}$ | SAFA/PUFA | >0.05 |
| $A\beta_{42}$/tau | C16:1/C16:0 | >0.05 |
| Tau | C16:1/C16:0 | >0.05 |
| $A\beta_{42}$ | C16:1/C16:0 | >0.05 |
| $A\beta_{42}$/tau | C20:2(n-6) | >0.05 |
| Tau | C20:2(n-6) | >0.05 |
| $A\beta_{42}$ | C20:2(n-6) | >0.05 |
| **CU vs ACD** | | |
| $A\beta_{42}$/tau | SDI | <0.01 |
| $A\beta_{42}$/tau | D9D | <0.01 |
| $A\beta_{42}$/tau | DI | <0.01 |
| $A\beta_{42}$/tau | C16:1 | <0.01 |
| $A\beta_{42}$/tau | C20:2(n-6) | <0.05 |
| $A\beta_{42}$/tau | C15:0 | <0.01 |
| $A\beta_{42}$/tau | N-3 | <0.05 |
| Tau | SDI | >0.05 |
| $A\beta_{42}$ | SDI | >0.05 |
| Tau | D9D | >0.05 |
| $A\beta_{42}$ | D9D | >0.05 |
| Tau | DI | >0.05 |
| Aβ42 | DI | >0.05 |
| Tau | C16:1 | >0.05 |
| $A\beta_{42}$ | C16:1 | >0.05 |
| Tau | C20:2(n-6) | >0.05 |
| $A\beta_{42}$ | C20:2(n-6) | >0.05 |
| Tau | C15:0 | >0.05 |
| $A\beta_{42}$ | C15:0 | >0.05 |
| Tau | N-3 | >0.05 |

| | | |
|---|---|---|
| Aβ42 | N-3 | >0.05 |

**Table S3: Multiple logistic regression of the best CSF unesterified fatty acids model for discriminating CU, MCI, and ACD**

| Model | N | AUC (95% CI) | Sensitivity | Specificity |
|---|---|---|---|---|
| **CU versus MCI** | | | | |
| Aβ42/tau + C17:0 | 68 vs 38 | 0.70(0.59-0.81) | 0.76 | 0.6 |
| Aβ42/tau + Aβ42 + C17:0 | 68 vs 38 | 0.70(0.59-0.81) | 0.74 | 0.63 |
| Aβ42/tau + Aβ42 + tau + C17:0 | 68 vs 38 | 0.70(0.59-0.80) | 0.74 | 0.65 |
| Aβ42/tau + Aβ42 + SAFA/PUFA | 68 vs 38 | 0.70(0.59-0.81) | 0.68 | 0.66 |
| Aβ42/tau + tau + SAFA/PUFA | 68 vs 38 | 0.70(0.59-0.80) | 0.53 | 0.79 |
| Aβ42/tau + SAFA/MUFA | 68 vs 38 | 0.70(0.59-0.80) | 0.55 | 0.78 |
| Aβ42/tau + Aβ42 + SAFA/MUFA | 68 vs 38 | 0.70(0.59-0.80) | 0.55 | 0.78 |
| Aβ42/tau + C16:1/C16:0 + SAFA/PUFA | 68 vs 38 | 0.70(0.59-0.80) | 0.4 | 0.94 |
| Aβ42/tau + C17:0 + C16:1/C16:0 + SAFA/PUFA | 68 vs 38 | 0.70(0.59-0.81) | 0.4 | 0.94 |
| **MCI versus ACD** | | | | |
| Aβ42/tau + D9D + DI | 38 vs 37 | 0.90(0.83-0.970 | 0.81 | 0.9 |
| Aβ42/tau + D9D + DI + SAFA/MUFA | 38 vs 37 | 0.90(0.83-0.97) | 0.73 | 0.95 |
| Aβ42/tau + SDI + DI | 38 vs 37 | 0.90(0.83-0.97) | 0.78 | 0.9 |
| Aβ42/tau + C20:2(n-6) + Aβ42 | 38 vs 37 | 0.89(0.81-0.97) | 0.92 | 0.79 |
| Aβ42/tau + C20:2(n-6) + D9D | 38 vs 37 | 0.89(0.80-0.97) | 0.92 | 0.79 |
| Aβ42/tau + C20:2(n-6) + DI | 38 vs 37 | 0.89(0.81-0.97) | 0.92 | 0.76 |
| Aβ42/tau + C20:2(n-6) + SAFA/MUFA | 38 vs 37 | 0.89(0.97) | 0.92 | 0.79 |
| Aβ42/tau + C20:2(n-6) | 38 vs 37 | 0.88(0.80-0.96) | 0.92 | 0.76 |
| Aβ42/tau + C20:2(n-6) + SAFA/PUFA | 38 vs 37 | 0.89(0.81-0.97) | 0.92 | 0.79 |
| Aβ42/tau + C20:2(n-6) + C16:1/C16:0 | 38 vs 37 | 0.89(0.81-0.97) | 0.92 | 0.79 |
| **CH versus ACD** | | | | |
| Aβ42/tau + C16:1 | 68 vs 37 | 0.87(0.80-0.95) | 0.78 | 0.84 |
| Aβ42/tau + C16:1 + Aβ42 | 68 vs 37 | 0.88(0.80-0.95) | 0.76 | 0.88 |
| Aβ42/tau + C16:1 + N-3 | 68 vs 37 | 0.88(0.81-0.95) | 0.78 | 0.85 |
| Aβ42/tau + C16:1 + C20:2(n-6) | 68 vs 37 | 0.87(0.80-0.95) | 0.76 | 0.88 |
| Aβ42/tau + DI + N-3 | 68 vs 37 | 0.87(0.80-0.95) | 0.87 | 0.77 |
| Aβ42/tau + DI + C16:1 | 68 vs 37 | 0.87(0.79-0.95) | 0.87 | 0.75 |
| Aβ42/tau + D9D + N-3 | 68 vs 37 | 0.87(0.79-0.95) | 0.78 | 0.87 |
| Aβ42/tau + SDI + N-3 | 68 vs 37 | 0.87(0.79-0.95) | 0.78 | 0.87 |
| Aβ42/tau + SDI + C16:1 | 68 vs 37 | 0.87(0.79-0.95) | 0.87 | 0.75 |

**Table S4: Best performing CSF supernatant fluid fatty acids for differential diagnosis of CU, MCI, and ACD based on ROC curve analyses.**

| Variable | N | AUC (95% CI) | Sensitivity | Specificity |
|---|---|---|---|---|
| **CU vs MCI** | | | | |
| C18:3(n-3) | 68 vs 38 | 0.66 (0.55-0.76) | 0.5 | 0.78 |
| N-3 | 68 vs 38 | 0.60 (0.49-0.72) | 0.71 | 0.55 |
| C16:0 | 68 vs 38 | 0.59 (0.48-0.70) | 0.55 | 0.67 |
| C20:4(n-6)/C22:6(n-3) | 68 vs 38 | 0.58 (0.46-0.69) | 0.61 | 0.63 |
| (C20:5(n-3)+C22:6(n-3))/C20:4(n-6) | 68 vs 38 | 0.57 (0.46-0.69) | 0.55 | 0.67 |
| C18:3(n-6) | 68 vs 38 | 0.56 (0.45-0.68) | 0.71 | 0.54 |
| MUFA/PUFA | 68 vs 38 | 0.56 (0.44-0.69) | 0.34 | 0.91 |
| C20:1 | 68 vs 38 | 0.56 (0.44-0.68) | 0.29 | 0.93 |
| C22:6(n-3) | 68 vs 38 | 0.56 (0.45-0.67) | 0.74 | 0.51 |
| PUFA | 68 vs 38 | 0.56 (0.44-0.67) | 0.58 | 0.6 |
| **MCI vs. ACD** | | | | |
| $A\beta_{42}$/tau | 38 vs 37 | 0.85 (0.75-0.94) | 0.89 | 0.76 |
| Tau | 38 vs 37 | 0.82 (0.72-0.92) | 0.73 | 0.84 |
| $A\beta_{42}$ | 38 vs 37 | 0.79 (0.68-0.89) | 0.78 | 0.82 |
| C20:1 | 38 vs 37 | 0.76 (0.64-0.87) | 0.73 | 0.71 |
| SDI | 38 vs 37 | 0.70 (0.58-0.82) | 0.73 | 0.66 |
| C20:3(n-3) | 38 vs 37 | 0.70 (0.58-0.82) | 0.62 | 0.76 |
| SAFA/MUFA | 38 vs 37 | 0.68 (0.55-0.80) | 0.78 | 0.61 |
| SAFA/PUFA | 38 vs 37 | 0.67 (0.55-0.80) | 0.81 | 0.55 |
| DI | 38 vs 37 | 0.67 (0.55-0.80) | 0.73 | 0.63 |
| **CU vs ACD** | | | | |
| $A\beta_{42}$/tau | 68 vs 37 | 0.85 (0.77-0.93) | 0.86 | 0.75 |
| Tau | 68 vs 37 | 0.80 (0.71-0.89) | 0.84 | 0.71 |
| $A\beta_{42}$ | 68 vs 37 | 0.76 (0.67-0.86) | 0.76 | 0.74 |
| C20:1 | 68 vs 37 | 0.73 (0.61-0.83) | 0.59 | 0.82 |
| N-3 | 68 vs 37 | 0.72 (0.63-0.82) | 0.89 | 0.45 |
| (C20:5(n-3)+C22:6(n-3))/C20:4(n-6) | 68 vs 37 | 0.71(0.61-0.82) | 0.68 | 0.7 |
| C20:4(n-6)/C22:6(n-3) | 68 vs 37 | 0.71(0.61-0.82) | 0.7 | 0.7 |
| C20:3(n-3) | 68 vs 37 | 0.70 (0.59-0.81) | 0.59 | 0.81 |
| C22:6(n-3) | 68 vs 37 | 0.68 (0.58-0.79) | 0.86 | 0.46 |
| SDI | 68 vs 37 | 0.66 (0.55-0.76) | 0.84 | 0.52 |

**Table S5: DeLong test for statistically significant difference between the AUC, sensitivity, and specificity of CSF supernatant fatty acids and Aβ42, tau, and Aβ42/tau**

| Predictor1 | Predictor2 | p-value |
|---|---|---|
| **CU vs MCI** | | |
| $A\beta_{42}$ | C18:3(n-3) | 0.04 |
| $A\beta_{42}$/tau | C18:3(n-3) | >0.05 |
| $A\beta_{42}$ | C18:3(n-3) | >0.05 |
| Tau | C18:3(n-3) | >0.05 |
| $A\beta_{42}$/tau | N-3 | >0.05 |
| $A\beta_{42}$ | N-3 | >0.05 |
| Tau | N-3 | >0.05 |
| $A\beta_{42}$/tau | C16:0 | >0.05 |
| $A\beta_{42}$ | C16:0 | >0.05 |
| Tau | C16:0 | >0.05 |
| $A\beta_{42}$/tau | C20:4n-6/C22:6n-3 | >0.05 |
| $A\beta_{42}$ | C20:4n-6/C22:6n-3 | >0.05 |
| Tau | C20:4n-6/C22:6n-3 | >0.05 |
| $A\beta_{42}$/tau | (C20:5n-3+C22:6n-3)/C20:4n-6 | >0.05 |
| $A\beta_{42}$ | (C20:5n-3+C22:6n-3)/C20:4n-6 | >0.05 |
| Tau | (C20:5n-3+C22:6n-3)/C20:4n-6 | >0.05 |
| $A\beta_{42}$/tau | C18:3(n-6) | >0.05 |
| $A\beta_{42}$ | C18:3(n-6) | >0.05 |
| Tau | C18:3(n-6) | >0.05 |
| $A\beta_{42}$/tau | MUFA/PUFA | >0.05 |
| $A\beta_{42}$ | MUFA/PUFA | >0.05 |
| Tau | MUFA/PUFA | >0.05 |
| $A\beta_{42}$/tau | C20:1 | >0.05 |
| $A\beta_{42}$ | C20:1 | >0.05 |
| Tau | C20:1 | >0.05 |
| $A\beta_{42}$/tau | C22:6(n-3) | >0.05 |
| $A\beta_{42}$ | C22:6(n-3) | >0.05 |
| Tau | C22:6(n-3) | >0.05 |
| $A\beta_{42}$/tau | PUFA | >0.05 |
| $A\beta_{42}$ | PUFA | >0.05 |
| Tau | PUFA | >0.05 |
| **MCI vs ACD** | | |
| $A\beta_{42}$/tau | DI | 0.03 |
| $A\beta_{42}$/tau | DI | 0.03 |
| $A\beta_{42}$/tau | SAFA/MUFA | 0.04 |
| $A\beta_{42}$/tau | SAFA/PUFA | 0.04 |

| | | |
|---|---|---|
| $A\beta_{42}$/tau | MUFA/PUFA | 0.002 |
| $A\beta_{42}$ | ecDI | >0.05 |
| Tau | ecDI | >0.05 |
| $A\beta_{42}$ | DI | >0.05 |
| Tau | DI | >0.05 |
| $A\beta_{42}$ | SAFA/MUFA | >0.05 |
| Tau | SAFA/MUFA | >0.05 |
| $A\beta_{42}$ | SAFA/PUFA | >0.05 |
| Tau | SAFA/PUFA | >0.05 |
| $A\beta_{42}$ | MUFA/PUFA | >0.05 |
| Tau | MUFA/PUFA | >0.05 |
| $A\beta_{42}$/tau | C20:1 | >0.05 |
| Tau | C20:1 | >0.05 |
| $A\beta_{42}$ | C20:1 | >0.05 |
| $A\beta_{42}$/tau | SDI | >0.05 |
| Tau | SDI | >0.05 |
| $A\beta_{42}$ | SDI | >0.05 |
| $A\beta_{42}$/tau | C20:3(n-3) | >0.05 |
| Tau | C20:3(n-3) | >0.05 |
| $A\beta_{42}$ | C20:3(n-3) | >0.05 |
| $A\beta_{42}$/tau | SAFA/MUFA | >0.05 |
| Tau | SAFA/MUFA | >0.05 |
| $A\beta_{42}$ | SAFA/MUFA | >0.05 |
| $A\beta_{42}$/tau | SAFA/PUFA | >0.05 |
| Tau | SAFA/PUFA | >0.05 |
| $A\beta_{42}$ | SAFA/PUFA | >0.05 |
| $A\beta_{42}$/tau | DI | >0.05 |
| Tau | DI | >0.05 |
| $A\beta_{42}$ | SF USFA/SAFA | >0.05 |
| $A\beta_{42}$/tau | SF USFA/SAFA | >0.05 |
| Tau | SF USFA/SAFA | >0.05 |
| $A\beta_{42}$ | SF USFA/SAFA | >0.05 |
| $A\beta_{42}$/tau | ecDI | >0.05 |
| Tau | ecDI | >0.05 |
| $A\beta_{42}$ | ecDI | >0.05 |
| | | >0.05 |
| **CU vs ACD** | | |
| Tau | SDI | 0.043 |
| $A\beta_{42}$/tau | SDI | 0.006 |
| $A\beta_{42}$/tau | C20:3(n-3) | 0.029 |
| $A\beta_{42}$/tau | C22:6(n-3) | 0.019 |

| | | |
|---|---|---|
| $A\beta_{42}$/tau | C20:4n-6/C22:6n-3 | 0.031 |
| $A\beta_{42}$/tau | (C20:5n-3+C22:6n-3)/C20:4n-6 | 0.032 |
| $A\beta_{42}$/tau | C20:1 | >0.05 |
| Tau | C20:1 | >0.05 |
| $A\beta_{42}$ | C20:1 | >0.05 |
| $A\beta_{42}$/tau | N-3 | >0.05 |
| Tau | N-3 | >0.05 |
| $A\beta_{42}$ | N-3 | >0.05 |
| $A\beta_{42}$/tau | (C20:5n-3+C22:6n-3)/C20:4n-6 | >0.05 |
| Tau | (C20:5n-3+C22:6n-3)/C20:4n-6 | >0.05 |
| $A\beta_{42}$ | (C20:5n-3+C22:6n-3)/C20:4n-6 | >0.05 |
| $A\beta_{42}$/tau | C20:4n-6/C22:6n-3 | >0.05 |
| Tau | C20:4n-6/C22:6n-3 | >0.05 |
| $A\beta_{42}$ | C20:4n-6/C22:6n-3 | >0.05 |
| $A\beta_{42}$/tau | C20:3(n-3) | >0.05 |
| Tau | C20:3(n-3) | >0.05 |
| $A\beta_{42}$ | C20:3(n-3) | >0.05 |
| $A\beta_{42}$/tau | C22:6(n-3) | >0.05 |
| Tau | C22:6(n-3) | >0.05 |
| $A\beta_{42}$ | C22:6(n-3) | >0.05 |
| $A\beta_{42}$ | SDI | >0.05 |

**Table S6: Multiple logistic regression of the best CSF supernatant fluid fatty acids model for discriminating CU, MCI, and ACD**

| Model | N | AUC (95% CI) | Sensitivity | Specificity |
|---|---|---|---|---|
| **CU vs MCI** | | | | |
| $A\beta_{42}$/tau + C18:3(n-3) | 68 vs 38 | 0.67(0.56-0.78) | 0.4 | 0.9 |
| $A\beta_{42}$/tau + tau + C18:3(n-3) | 68 vs 38 | 0.67(0.56-0.78) | 0.42 | 0.87 |
| $A\beta_{42}$/tau + $A\beta_{42}$ + C18:3(n-3) | 68 vs 38 | 0.66(0.56-0.77) | 0.82 | 0.46 |
| $A\beta_{42}$/tau + N-3 | 68 vs 38 | 0.61(0.50-0.72) | 0.66 | 0.55 |
| $A\beta_{42}$/tau + tau + N-3 | 68 vs 38 | 0.62(0.51-0.73) | 0.74 | 0.52 |
| $A\beta_{42}$/tau + $A\beta_{42}$ + N-3 | 68 vs 38 | 0.61(0.49-0.72) | 0.68 | 0.52 |
| $A\beta_{42}$/tau + C18:3(n-3) + N-3 | 68 vs 38 | 0.67(0.56-0.78) | 0.45 | 0.82 |
| $A\beta_{42}$/tau + C16:0 | 68 vs 38 | 0.61(0.50-0.72) | 0.47 | 0.73 |
| $A\beta_{42}$/tau + (C20:5n-3+C22:6n-3)/C20:4n-6 | 68 vs 38 | 0.60(0.49-0.72) | 0.55 | 0.66 |
| **MCI vs ACD** | | | | |
| $A\beta_{42}$/tau + C20:3(n-3) + C20:1 | 38 vs 37 | 0.94(0.89-0.94) | 0.97 | 0.82 |
| $A\beta_{42}$/tau + C20:1 + N-3 + $A\beta_{42}$ | 38 vs 37 | 0.94(0.88-0.94) | 0.92 | 0.9 |

| | | | | |
|---|---|---|---|---|
| $A\beta_{42}$/tau + C20:1 + N-3 | 38 vs 37 | 0.93(0.87-0.93) | 0.95 | 0.82 |
| $A\beta_{42}$/tau + tau + SDI | 38 vs 37 | 0.93(0.86-0.93) | 0.89 | 0.87 |
| $A\beta_{42}$/tau + C20:1 + SDI | 38 vs 37 | 0.93(0.86-0.93) | 0.89 | 0.87 |
| $A\beta_{42}$/tau + $A\beta_{42}$ + C20:1 | 38 vs 37 | 0.92(0.85-0.92) | 0.97 | 0.79 |
| $A\beta_{42}$/tau + C20:1 | 38 vs 37 | 0.92(0.85-0.92) | 0.97 | 0.79 |
| $A\beta_{42}$/tau + C20:1 + tau | 38 vs 37 | 0.92(0.85-0.92) | 0.94 | 0.81 |
| $A\beta_{42}$/tau + SDI | 38 vs 37 | 0.90(0.83-90.0) | 0.86 | 0.92 |
| **CU vs ACD** | | | | |
| $A\beta_{42}$/tau + C20:1 | 68 vs 37 | 0.88(0.80-0.95) | 0.78 | 0.84 |
| $A\beta_{42}$/tau + tau + C20:1 | 68 vs 37 | 0.87(0.80-95) | 0.76 | 0.87 |
| $A\beta_{42}$/tau + $A\beta_{42}$ + C20:1 | 68 vs 37 | 0.88(0.80-0.95) | 0.78 | 0.84 |
| $A\beta_{42}$/tau + N-3 | 68 vs 37 | 0.88(0.81-0.95) | 0.87 | 0.78 |
| $A\beta_{42}$/tau + tau + N-3 | 68 vs 37 | 0.88(0.82-0.95) | 0.87 | 0.79 |
| $A\beta_{42}$/tau + $A\beta_{42}$ + N-3 | 68 vs 37 | 0.89(0.82-95) | 0.78 | 0.88 |
| $A\beta_{42}$/tau + C20:1 + N-3 | 68 vs 37 | 0.92(0.86-0.98) | 0.81 | 0.91 |
| $A\beta_{42}$/tau + $A\beta_{42}$ + C20:1 + N-3 | 68 vs 37 | 0.92(0.86-0.98) | 0.81 | 0.93 |
| $A\beta_{42}$/tau + tau + C20:1 + N-3 | 68 vs 37 | 0.92(0.86-0.98) | 0.92 | 0.82 |

**Table S7: Best performing CSF nanoparticles fatty acids for differential diagnosis of CU, MCI, and ACD based on ROC curve analyses**

| Variable | N | AUC (95% CI) | Sensitivity | Specificity |
|---|---|---|---|---|
| **CU vs MCI** | | | | |
| C16:1 | 68 vs 38 | 0.71(0.61-0.81) | 0.53 | 0.81 |
| C17:0 | 68 vs 38 | 0.69(0.58-0.79) | 0.5 | 0.84 |
| C15:0 | 68 vs 38 | 0.69(0.58-0.79) | 0.71 | 0.62 |
| C15:1/C15:0 | 68 vs 38 | 0.69(0.58-0.79) | 0.58 | 0.76 |
| C14:0/C16:0 | 68 vs 38 | 0.69(0.58-0.79) | 0.47 | 0.84 |
| ocSAFA | 68 vs 38 | 0.69(0.58-0.79) | 0.82 | 0.53 |
| (C14:0+C16:0)/(C16:0+C18:0) | 68 vs 38 | 0.68(0.57-0.78) | 0.63 | 0.71 |
| C16:0/C18:0 | 68 vs 38 | 0.67(0.56-0.78) | 0.63 | 0.68 |
| C14:1 | 68 vs 38 | 0.67(0.56-0.77) | 0.76 | 0.57 |
| ocDI | 68 vs 38 | 0.67(0.56-0.77) | 0.55 | 0.74 |
| **MCI vs ACD** | | | | |
| $A\beta_{42}$/tau | 38 vs 37 | 0.85(0.75-0.94) | 0.89 | 0.76 |
| Tau | 38 vs 37 | 0.82(0.72-0.92) | 0.73 | 0.84 |

| | | | | |
|---|---|---|---|---|
| Aβ42 | 38 vs 37 | 0.79(0.68-0.89) | 0.78 | 0.82 |
| C15:1/C15:0 | 38 vs 37 | 0.75(0.64-0.86) | 0.89 | 0.53 |
| C14:0/C16:0 | 38 vs 37 | 0.72(0.61-0.84) | 0.92 | 0.53 |
| ocDI | 38 vs 37 | 0.72(0.61-0.84) | 0.81 | 0.55 |
| C20:5(n-3) | 38 vs 37 | 0.72(0.60-0.83) | 0.67 | 0.74 |
| ecDI | 38 vs 37 | 0.71(0.59-0.83) | 0.72 | 0.71 |
| DI | 38 vs 37 | 0.71(0.59-0.83) | 0.72 | 0.68 |
| SAFA/MUFA | 38 vs 37 | 0.71(0.59-0.83) | 0.75 | 0.63 |
| **CU vs ACD** | | | | |
| Aβ42/tau | 68 vs 37 | 0.85(0.77-0.93) | 0.87 | 0.75 |
| Tau | 68 vs 37 | 0.80(0.72-0.89) | 0.84 | 0.71 |
| Aβ42 | 68 vs 37 | 0.76(0.67-0.86) | 0.76 | 0.74 |
| C20:5(n-3) | 68 vs 37 | 0.65(0.54-0.76) | 0.53 | 0.75 |
| C15:1 | 68 vs 37 | 0.62(0.51-0.74) | 0.61 | 0.66 |
| ecDI | 68 vs 37 | 0.62(0.50-0.73) | 0.72 | 0.54 |
| ocMUFA | 68 vs 37 | 0.61(0.49-0.73) | 0.61 | 0.65 |
| C20:3(n-3) | 68 vs 37 | 0.61(0.50-0.73) | 0.78 | 0.47 |
| SAFA/MUFA | 68 vs 37 | 0.61(0.50-0.73) | 0.69 | 0.54 |
| DI | 68 vs 37 | 0.61(0.50-0.73) | 0.72 | 0.52 |

**Table S8: DeLong test for statistically significant difference between the AUC, sensitivity, and specificity of CSF nanoparticles fatty acids and Aβ42, tau, and Aβ42/tau**

| Predictor1 | Predictor2 | p-value |
|---|---|---|
| **CU vs MCI** | | |
| Aβ42 | C15:0 | 0.045 |
| Aβ42 | C16:1 | 0.019 |
| Aβ42 | C15:1/C15:0 | 0.042 |
| Aβ42 | C17:0 | 0.041 |
| Aβ42 | ocSAFA | 0.043 |
| Aβ42 | C14:0/C16:0 | 0.037 |
| Tau | C16:1 | 0.019 |
| Aβ42/tau | C16:1 | 0.035 |
| Tau | C15:0 | >0.05 |
| Aβ42/tau | C15:0 | >0.05 |
| Tau | C16:1 | >0.05 |
| Tau | C15:1/C15:0 | >0.05 |
| Aβ42/tau | C15:1/C15:0 | >0.05 |
| Tau | C17:0 | >0.05 |
| Aβ42/tau | C17:0 | >0.05 |

| | | |
|---|---|---|
| Tau | ocSAFA | >0.05 |
| $A\beta_{42}$/tau | ocSAFA | >0.05 |
| Tau | C14:0/C16:0 | >0.05 |
| $A\beta_{42}$/tau | C14:0/C16:0 | >0.05 |
| **MCI vs ACD** | | |
| $A\beta_{42}$/tau | SAFA/MUFA | 0.044 |
| $A\beta_{42}$/tau | SAFA/MUFA | 0.044 |
| $A\beta_{42}$/tau | C15:1/C15:0 | >0.05 |
| $A\beta_{42}$ | C15:1/C15:0 | >0.05 |
| Tau | C15:1/C15:0 | >0.05 |
| $A\beta_{42}$/tau | C14:0/C16:0 | >0.05 |
| $A\beta_{42}$ | C14:0/C16:0 | >0.05 |
| Tau | C14:0/C16:0 | >0.05 |
| $A\beta_{42}$/tau | ocDI | >0.05 |
| $A\beta_{42}$ | ocDI | >0.05 |
| Tau | ocDI | >0.05 |
| $A\beta_{42}$/tau | C20:5(n-3) | >0.05 |
| $A\beta_{42}$ | C20:5(n-3) | >0.05 |
| Tau | C20:5(n-3) | >0.05 |
| $A\beta_{42}$/tau | ecDI | >0.05 |
| $A\beta_{42}$ | ecDI | >0.05 |
| Tau | ecDI | >0.05 |
| $A\beta_{42}$/tau | SAFA/MUFA | >0.05 |
| $A\beta_{42}$ | SAFA/MUFA | >0.05 |
| Tau | SAFA/MUFA | >0.05 |
| **CU vs ACD** | | |
| $A\beta_{42}$ | C15:1 | 0.044 |
| $A\beta_{42}$ | DI | 0.031 |
| $A\beta_{42}$ | ecDI | 0.038 |
| $A\beta_{42}$ | C20:3(n-3) | 0.034 |
| $A\beta_{42}$ | ocMUFA | 0.037 |
| $A\beta_{42}$ | SAFA/MUFA | 0.033 |
| Tau | C15:1 | 0.009 |
| Tau | DI | 0.01 |
| Tau | ecDI | 0.012 |
| Tau | C20:5(n-3) | 0.033 |
| Tau | C20:5(n-3) | 0.006 |
| Tau | SAFA/MUFA | 0.01 |
| Tau | ocMUFA | 0.007 |
| $A\beta_{42}$/tau | C15:1 | 0 |
| $A\beta_{42}$/tau | ecDI | 0.001 |

| | | |
|---|---|---|
| $A\beta_{42}$/tau | DI | 0 |
| $A\beta_{42}$/tau | C20:5(n-3) | 0.004 |
| $A\beta_{42}$/tau | C20:3(n-3) | 0 |
| $A\beta_{42}$/tau | MUFA | 0 |
| $A\beta_{42}$/tau | SAFA/MUFA | 0.001 |

**Table S9: Multiple logistic regression of the best CSF nanoparticles fatty acids model for discriminating CU, MCI, and ACD**

| Variable | N | AUC (95% CI) | Sensitivity | Specificity |
|---|---|---|---|---|
| **CU vs MCI** | | | | |
| $A\beta_{42}$/tau + C16:1 | 68 vs 38 | 0.71(0.61-0.81) | 0.76 | 0.65 |
| $A\beta_{42}$/tau + C16:1 + $A\beta_{42}$ | 68 vs 38 | 0.71(0.61-0.81) | 0.71 | 0.66 |
| $A\beta_{42}$/tau + C16:1 + tau | 68 vs 38 | 0.71(0.61-0.81) | 0.74 | 0.62 |
| $A\beta_{42}$/tau + C17:0 | 68 vs 38 | 0.69(0.59-0.80) | 0.84 | 0.52 |
| $A\beta_{42}$/tau + C17:0 + $A\beta_{42}$ | 68 vs 38 | 0.69(0.59-0.80) | 0.84 | 0.5 |
| $A\beta_{42}$/tau + C17:0 + tau | 68 vs 38 | 0.69(0.58-0.79) | 0.92 | 0.43 |
| $A\beta_{42}$/tau + C15:0 | 68 vs 38 | 0.69(0.58-0.80) | 0.79 | 0.54 |
| **MCI vs. ACD** | | | | |
| $A\beta_{42}$/tau + C15:1/C15:0 | 38 vs 37 | 0.88(0.80-0.97) | 0.81 | 0.9 |
| $A\beta_{42}$/tau + tau + C15:1/C15:0 | 38 vs 37 | 0.88(0.80-0.97) | 0.81 | 0.92 |
| $A\beta_{42}$/tau + $A\beta_{42}$ + C15:1/C15:0 | 38 vs 37 | 0.88(0.79-0.97) | 0.81 | 0.92 |
| $A\beta_{42}$/tau + C15:1/C15:0 + C14:0/C16:0 | 38 vs 37 | 0.88(0.80-0.97) | 0.81 | 0.9 |
| $A\beta_{42}$/tau + C14:0/C16:0 | 38 vs 37 | 0.89(0.80-0.97) | 0.78 | 0.92 |
| $A\beta_{42}$/tau + $A\beta_{42}$ + C14:0/C16:0 | 38 vs 37 | 0.88(0.80-0.97) | 0.94 | 0.74 |
| $A\beta_{42}$/tau + tau + C14:0/C16:0 | 38 vs 37 | 0.89(0.82-0.97) | 0.92 | 0.76 |
| $A\beta_{42}$/tau + C20:5(n-3) | 38 vs 37 | 0.89(0.82-0.97) | 0.92 | 0.74 |
| $A\beta_{42}$/tau + C20:5(n-3) + $A\beta_{42}$ | 38 vs 37 | 0.88(0.80-0.96) | 0.81 | 0.87 |
| $A\beta_{42}$/tau + C20:5(n-3) + tau | 38 vs 37 | 0.89(0.81-0.97) | 0.92 | 0.74 |
| $A\beta_{42}$/tau + ocDI | 38 vs 37 | 0.88(0.80-0.96) | 0.81 | 0.87 |
| **CU vs ACD** | | | | |
| $A\beta_{42}$/tau + C20:5(n-3) + C15:1 + tau | 68 vs 37 | 0.86(0.79-0.94) | 0.86 | 0.75 |
| $A\beta_{42}$/tau + C20:5(n-3) | 68 vs 37 | 0.85(0.77-0.92) | 0.72 | 0.84 |
| $A\beta_{42}$/tau + C20:5(n-3) + $A\beta_{42}$ | 68 vs 37 | 0.84(0.77-0.92) | 0.64 | 0.91 |
| $A\beta_{42}$/tau + C20:5(n-3) + tau | 68 vs 37 | 0.85(0.77-0.93) | 0.89 | 0.72 |

| | | | | |
|---|---|---|---|---|
| $A\beta_{42}$/tau + C20:5(n-3) + C15:1 | 68 vs 37 | 0.86(0.78-0.93) | 0.69 | 0.9 |
| $A\beta_{42}$/tau + C20:5(n-3) + C15:1 + $A\beta_{42}$ | 68 vs 37 | 0.86(0.78-0.93) | 0.69 | 0.9 |
| $A\beta_{42}$/tau + tau | 68 vs 37 | 0.85(0.77-0.93) | 0.89 | 0.78 |
| $A\beta_{42}$/tau + $A\beta_{42}$ | 68 vs 37 | 0.85(0.77-0.94) | 0.89 | 0.75 |
